# Bacterial metabolism of synthetic steroids across ecosystems reveals diverse biotransformation products, reactions, and enzymes

**DOI:** 10.64898/2026.06.05.730323

**Authors:** Mariia A. Beliaeva, Matthias W. Gross, Dimitrios Papagiannidis, Anoop Singh, Mikhail Savitski, Michael Zimmermann

## Abstract

Biotransformation of synthetic steroid drugs by gut and environmental bacteria affects their activity within the host, persistence, and environmental fate of these widely used pharmaceuticals. However, the spectrum of bacterial transformation of synthetic steroids, the enzymes involved, and the role of cooperative microbial metabolism in these processes remain poorly understood. We systematically mapped the biotransformation of 22 steroids, comprising 20 synthetic and 2 natural compounds and spanning clinically used estrogens, progestogens, corticosteroids, and prodrugs, across 12 bacterial species (8 intestinal and 4 environmental), yielding 264 steroid-bacteria combinations. We identified 97 unique biotransformation products and found that the tested bacteria catalyze diverse reactions including ester hydrolysis, oxidation-reduction chemistry, and steroid side-chain cleavage. Notably, the environmental bacterium *Sphingobium herbicidovorans* catalyzed desmolase-like steroid side-chain cleavage under aerobic conditions, a transformation previously reported solely for anaerobic bacteria. Combining sequence homology searches, a genetic gain-of-function screen, and expression proteomics we identified six steroid-transforming enzymes in *S. herbicidovorans*. We further demonstrate that distinct bacterial species cooperatively metabolize synthetic steroids through cross-feeding, enabling sequential activation and metabolism of corticosteroids across microbial communities. Together, our findings uncover previously unrecognized bacterial enzymes and community-level interactions involved in synthetic steroid metabolism. By directly linking steroid metabolites, enzymes, and microbial community interactions, this study provides molecular-level mechanistic insights into microbial steroid biotransformation. Such insights are essential for ultimately predicting metabolic interactions within and across microbial communities, as well as their interactions with the host and the environment.

## INTRODUCTION

Steroid drugs are essential medicines used to treat a variety of diseases, with three major classes corticosteroids, progestogens, and estrogens accounting for 12% of the most frequently prescribed drugs (clincalc.com) (Fig. 1a). In fact, many of the commercial steroid drugs are synthetic analogs of natural hormones, such as the anti-inflammatory drug dexamethasone and the contraceptives levonorgestrel and 17α-ethinylestradiol (EE2). Besides human medication, steroids are also commonly used in veterinary practices (*i.e.*, the estrus suppression drug altrenogest). To increase drug potency and avoid excessive host metabolism, most synthetic steroids contain artificial chemical moieties, such as ethynyl groups, the halogens fluorine and chlorine, which grant them greater metabolic stability compared to natural steroid hormones. However, introduction of such functional groups also leads to increased resilience of synthetic hormones to environmental degradation, resulting in their accumulation and associated negative effects on ecosystems^1^. For example, even sub-nanomolar concentration of artificial steroids in natural waters have been shown to disrupt the development and reproduction of fish and amphibians^2–6^ and pose the potential risks to plant growth^7^.

**Fig. 1.**
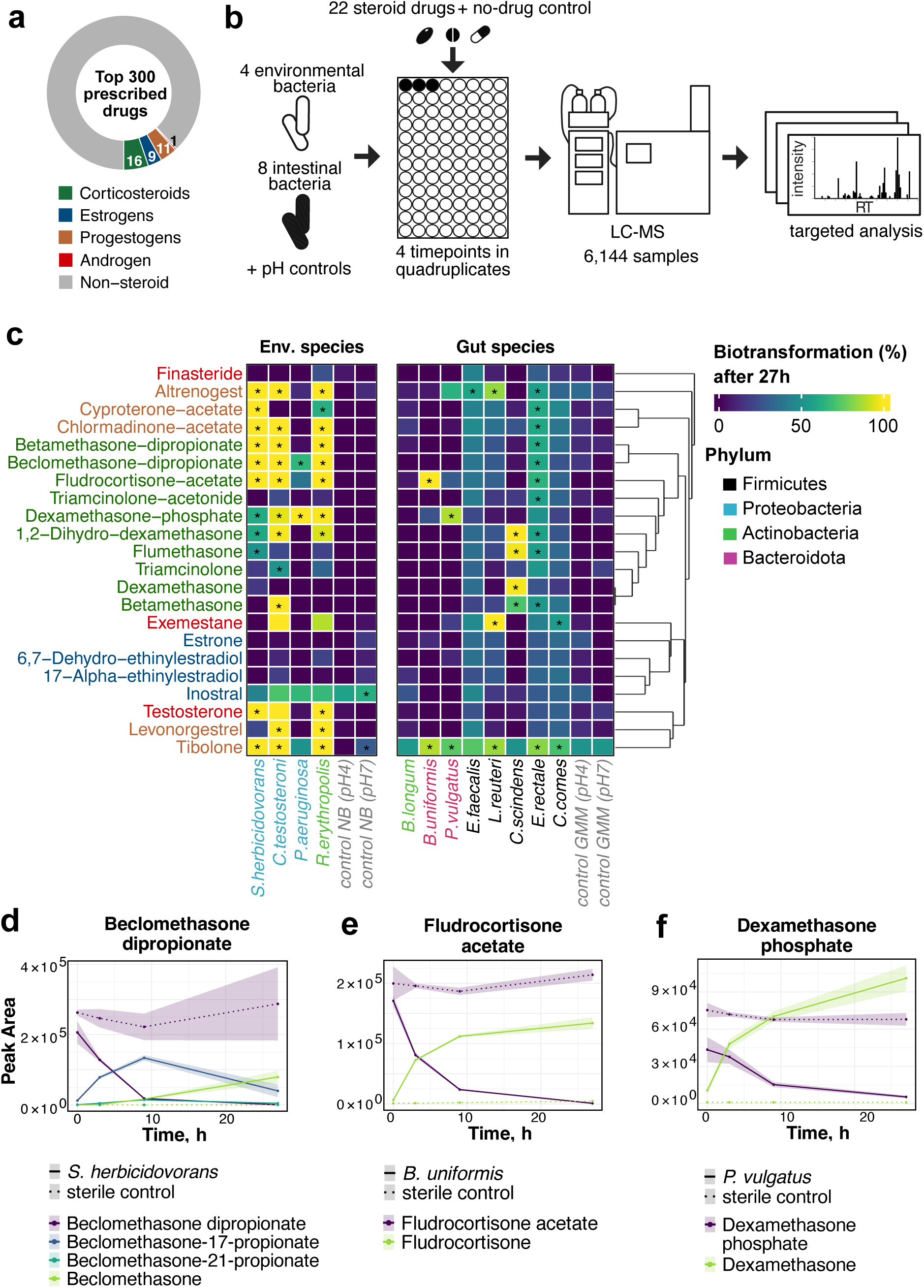
Assay to probe bacterial biotransformation of synthetic steroids. **a,** Share of drugs with steroidal scaffold out of the 300 most prescribed pharmaceuticals in 2023 according to ClinCalc DrugStats Database (clincalc.com/DrugStats/Top300Drugs.aspx). **b,** Scheme of the steroid biotransformation assay. **c,** Heatmap of the steroid–bacteria metabolism (n = 4 replicates); interactions with significant steroid transformation (≥ 50%, two-way ANOVA, FDR-corrected p-value ≤ 0.05, Benjamini-Hochberg) are marked with an asterisk. Steroids are arranged by their chemical similarity based on Tanimoto distances, and strains are arranged by their taxonomic relationships. **d-f,** Time-course of two-step hydrolysis of beclomethasone dipropionate by *S. herbicidovorans* (d), hydrolysis of fludrocortisone acetate to fludrocortisone by *B. uniformis* (e) and hydrolysis of dexamethasone phosphate to dexamethasone by *P. vulgatus* (f) (n = 4 replicates; shaded area, mean ± SD).

Microbial metabolism of synthetic steroids has been largely studied in two separate contexts: mineralization or degradation by environmental bacteria, and biotransformation by host-associated bacteria. In environmental microbiology, studies of steroid drugs have primarily focused on oxidative degradation and de-functionalization reactions associated with steroid catabolism. For example, metabolism of 17α-ethinylestradiol by a few environmental species has been reported, although degradation proceeds substantially slower than for natural analogues^8–11^. Similarly, activated-sludge studies demonstrated inefficient removal of halogenated corticosteroids and steroids containing C16,C17-cyclic ketal^12–14^, and partial degradation of dexamethasone through A-ring cleavage was reported in *Pseudomonas alcaligenes* and *Rhodococcus sp. D32*^15, 16^. Defluorination of triamcinolone acetonide by *Acinetobacter pittii* C3 has likewise been described, although it requires prolonged incubation^17, 18^. In contrast to these studies, which predominantly examine steroid degradation, investigations of host-associated bacteria have uncovered enzymes that selectively remodel steroid structures without necessarily supporting mineralization but influencing bioavailability and host physiology. Human gut bacteria have been reported to catalyze a distinct set of biotransformation reactions including dehydroxylation, dehydrogenation, side-chain cleavage, and reduction and oxidations, providing mechanistic insights into steroid core modifications^19–21^. For example, *Bacteroides thetaiotaomicron* encodes a 5α-reductase capable of transforming norethindrone and several additional steroids^22, 23^, whereas *Clostridium scindens* converts corticosteroids into androgenic 17-ketosteroids through desmolase activity^24^. More recently, the genes responsible for the transformation of corticoids into progestins through 21-hydroxylation were identified in *Eggerthella lenta*^25^. In addition, the gut bacterium *Clostridium steroidoreducens* was shown to reduce cortisol and related natural and synthetic steroid hormones to 3β,5β-tetrahydrosteroid products through the reductive OsrABC pathway, in which the 3-oxo-Δ¹-steroid hormone reductase OsrA acts on synthetic glucocorticoids such as prednisolone^26^. Taken together, these studies demonstrate that bacteria in various environments can catalyze chemically diverse transformations of steroids, potentially altering their biological activity, toxicity, persistence, and interactions with host steroid metabolism. However, current knowledge remains limited to isolated examples of individual studies, while a broader understanding of the chemical diversity of bacterial steroid biotransformation, the involved enzymes, and the contribution of microbial communities and their interplay to these processes is mostly lacking. We systematically profiled the biotransformation of 22 steroids across 4 environmental and 8 intestinal bacterial species using an MS/MS-based analytical pipeline to identify transformation products and resolve underlying metabolic activities. In total, we mapped 264 steroid-bacteria interactions and identified 97 unique steroid metabolites, revealing a broad spectrum of reactions including hydrolysis, oxidation, reduction, and C-C bond cleavage across diverse steroid scaffolds (spanning synthetic estrogens, progestogens, corticosteroids and selected prodrugs together with the natural steroids testosterone and estrone). Further, we combined bioinformatic predictions, a genetic gain-of-function screen, and expression proteomics to identify six enzymes in the environmental bacterium *Sphingobium herbicidovorans*. These enzymes are responsible for distinct steroid-metabolizing reactions, such as aerobic desmolase activity previously thought to be restricted to anaerobic gut bacteria. Finally, we employed cross-feeding experiments with axenic and microbial community cultures to demonstrate that environmental and intestinal bacteria can act cooperatively to metabolize steroids.

Taken together, our findings show that bacterial steroid biotransformation is more diverse than previously assumed and can be highly organism- and environment-specific. At the same time, steroid metabolism can arise from a combination of specialized enzymatic activities and cooperative processes within microbial communities. Resolving the enzymatic steps underlying these transformations is therefore essential for predicting the metabolic fate of synthetic steroids within and across microbial ecosystems. In addition, identifying these activities and their associated enzymes provides opportunities to harness them for biotechnological and bioremediation applications targeting steroid-based compounds.

## RESULTS

### Synthetic steroid biotransformation reactions by gut and environmental bacteria

To systematically assess the breadth and specificity of microbial steroid metabolism across environmental and gut-associated bacteria, we performed a biotransformation screen using a chemically diverse panel of steroid compounds. We assessed biotransformation of 20 synthetic steroid drugs, spanning estrogens, progestogens and corticosteroids, and respective prodrugs, and the natural steroids testosterone and estrone (Supplementary Table S1) by four environmental and eight intestinal bacterial species of four distinct phyla. Species were selected based on either reported metabolism of steroid compounds, such as hormones, bile acids or drugs (*C. scindens*^24^, *Comamonas testosteroni*^27^, *Rhodococcus erythropolis*^28^*, Pseudomonas aeruginosa*^29^*, Enterococcus faecalis*^30^,), or reported association with steroid hormones and health (*Bifidobacterium longum subsp. infantis*^31^*, Coprococcus comes, Eubacterium rectale*^32^*, Limosilactobacillus reuteri*^32^ (former *Lactobacillus reuteri*)) or overall ability to biotransform xenobiotics (*Sphingobium herbicidovorans*^33^*, Bacteroides uniformis*^22^*, Phocaeicola vulgatus*^34^) (Supplementary Table S2). Each compound was individually incubated at 5 μM with axenic bacterial cultures in quadruplicate, and samples were collected at 0, 3, 9, and 27 h of incubation under aerobic and anaerobic conditions for environmental and intestinal bacteria, respectively. In total, 6,144 samples including bacterial assays and sterile controls at pH 4 and pH 7 were measured by liquid-chromatography-coupled mass spectrometry (LC-MS) (Fig. 1b), operating at high mass resolution in scanning mode. We employed targeted data analysis to monitor the levels of parent compounds. Based on this analysis, we found steroid biotransformation (≥50% depletion of the parent compound, two-way ANOVA, FDR-corrected p-value ≤ 0.05, Benjamini-Hochberg, n = 4) in 35 of 88 environmental and 24 of 176 gut bacteria-steroid interactions after 27 h incubation (Fig. 1c). 17 of 22 tested steroids were metabolized by at least one bacterial species, and the five compounds that were not significantly transformed by any of the tested strains are the aza-steroid finasteride and the estrogens estrone, 17α-ethinylestradiol, 6,7-dehydroethinylestradiol and inostral.

In contrast, multiple corticosteroids and their esterified prodrugs showed pronounced biotransformation, particularly among environmental species. Among the gut bacterial species, *C. scindens* exhibited the strongest biotransformation activity, converting the corticosteroids flumethasone, dexamethasone, and betamethasone into their respective 17-ketosteroids (Supplementary Fig. 1a). In contrast, biotransformation of triamcinolone, which differs from dexamethasone only by a 16-hydroxyl group replacing a methyl group, was minimal, highlighting the structural specificity of the previously reported desmolase activity of *C. scindens*^24^. All prodrugs included in the screen (*i.e.*, fludrocortisone acetate, betamethasone dipropionate, beclomethasone dipropionate, and dexamethasone phosphate) were metabolized by at least one bacterial species, suggesting microbial activation of these compounds. To explicitly test this hypothesis, we performed suspect screening for the corresponding active drug molecules (Supplementary Table S4) and confirmed prodrug activation in all cases. For example, *S. herbicidovorans* converted beclomethasone dipropionate to beclomethasone *via* stepwise hydrolysis of the propionyl moieties, demonstrating slightly different kinetics for 17-propionate and 21-propionate formation, which is likely due to sterically hindered accessibility of the ester bond in position 17 (Fig. 1d, Supplementary Fig. 1b). Among gut-associated species, prodrug hydrolysis was more limited and was observed only for fludrocortisone acetate activated by *B. uniformis* (Fig. 1e, Supplementary Fig. 1c) and dexamethasone phosphate activated by *P. vulgatus* (Fig. 1f, Supplementary Fig. 1d). This targeted data analysis is inherently limited to predefined biotransformation hypotheses and cannot identify new steroid metabolites and new biotransformation reactions and hence fails to capture the full diversity of biotransformation. This limitation motivated us to apply a more agnostic strategy to comprehensively identify bacterial biotransformation products.

### Untargeted metabolomics and MS/MS analysis systematically identifies steroid metabolites

To systematically identify steroid biotransformation products, we developed a pipeline consisting of the following three steps (Fig. 2a): i) untargeted metabolomics followed by differential analysis to identify putative steroid metabolites; ii) targeted acquisition of the MS/MS spectra for steroids and their putative metabolites; iii) spectral similarity analysis between steroids and their putative metabolites. We selected 74 steroid-bacteria combinations (37 with environmental and 37 with gut bacteria) for untargeted identification of putative biotransformation products (Supplementary Table S5): 68 combinations reaching ≥50% steroid depletion at one or more timepoints relative to the paired abiotic controls, together with 6 combinations below this threshold that were retained to probe for low-level metabolite formation. We subjected the respective samples of the initial screen to untargeted metabolomics data analysis, resulting in >2,000 metabolic features for each of 74 steroid-bacteria pairs. To select molecular features for MS/MS analysis, we performed a differential abundance analysis across the time course for each steroid-bacteria combination. Features were prioritized through the following filters: (i) time-resolved differential abundance, retaining features that changed between later timepoints (3, 9, or 27 h) and 0 h with a fold change (FC, expressed in log2) ≥ 1 and FDR-corrected p-value ≤ 0.05 (Benjamini-Hochberg), based on quadruplicates (n = 4); (ii) removal of features also detected in the no-drug (DMSO) control, to exclude produced metabolites independent of the steroid present. Features recurring across timepoints within a given steroid-bacteria combination were counted once. This yielded 9,971 putative metabolites across all 74 steroid-bacteria pairs (Supplementary Table S6). For each steroid-bacteria pair, the corresponding ion list (23 to 218 features), together with the ions of the original steroid where these had not already been captured, was added to the inclusion list for subsequent MS/MS acquisition, giving 296 MS/MS acquisition runs in total. Chemical standards of steroids were used to optimize the collision energy (CE) to 10 eV, as it resulted in both well-defined precursor ions and informative fragment ions, whereas 5, 20, and 40 eV gave less optimal results (Supplementary Fig. 2a).

**Fig. 2.**
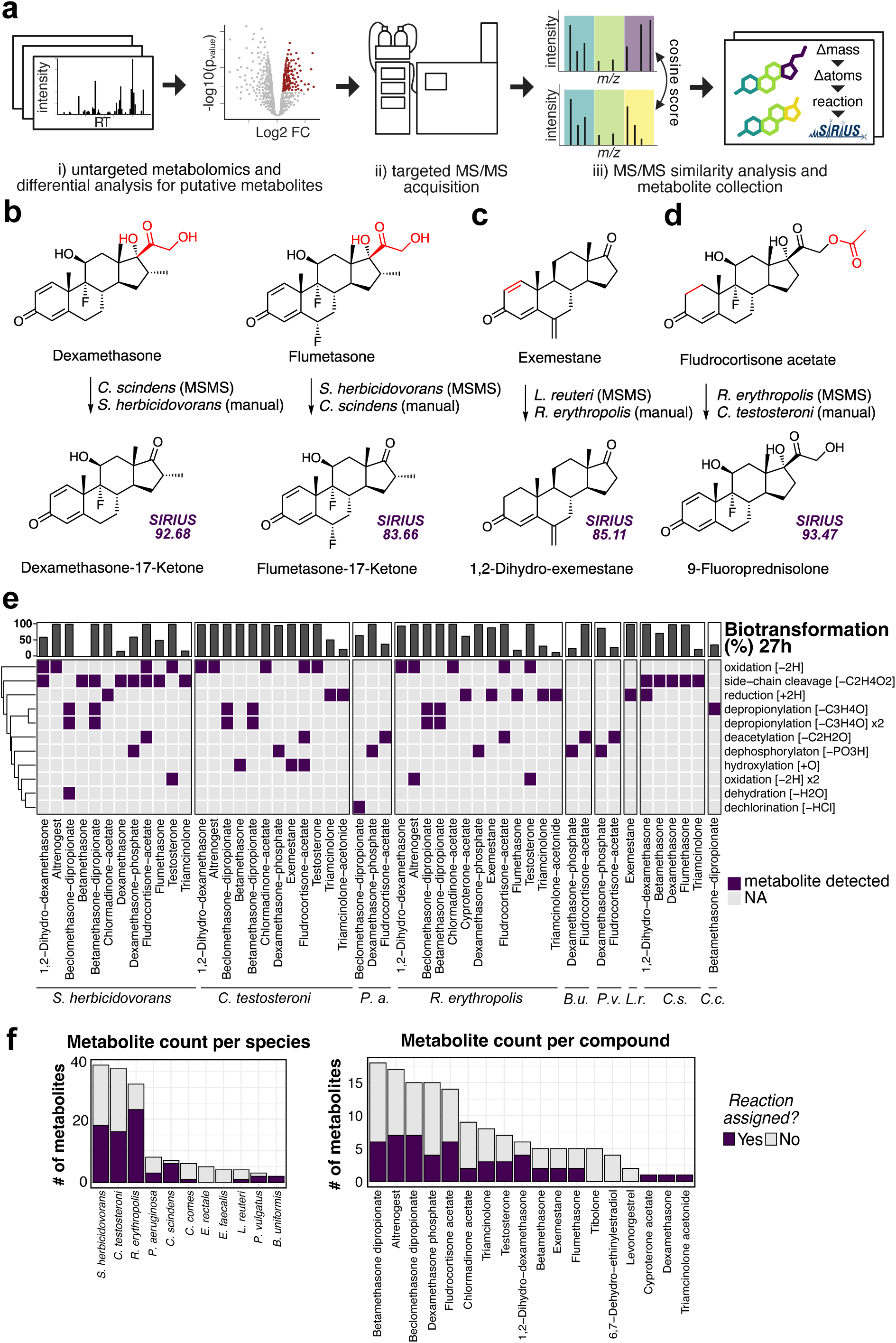
Identification of steroid metabolites. **a,** Scheme of the MS/MS pipeline to identify steroid biotransformation products. **b-d,** Examples of biotransformation reactions assigned to specific steroid-bacteria interactions: side-chain cleavage of dexamethasone and flumethasone by *C. scindens* and *S. herbicidovorans* resulting in respective 17-ketosteroids (b), reduction of exemestane to 1,2-dihydro-exemestane by gut bacteria *L. reuteri* and environmental bacteria *R. erythropolis* (c), biotransformation of fludrocortisone acetate by environmental species *R. erythropolis* and *C. testosteroni* depicting acetate hydrolysis and oxidation of A-ring (d). **e,** Full map of the biotransformation reactions that result in metabolites identified by MS/MS and subsequent suspect-screening. **f,** Number of metabolites observed per species and per steroid and the portion of metabolites with assigned reactions.

Next, we generated MS/MS spectral libraries for each steroid-bacteria combination by combining spectra across all timepoints, and used the steroid drugs and known prodrug metabolites (Supplementary Table S4) as references to compute spectral similarity with the cosine and F1 algorithms to leverage their complementary strengths and maximize capturing of structurally related metabolites^35^. We then applied the following filters to short-list putative metabolites (Supplementary Fig. 2, b): (i) spectral similarity to the reference spectra, retaining features above either threshold (cosine ≥ 0.5 or F1 ≥ 0.1); (ii) a time-resolved abundance change within the bacterial culture, retaining features with a significant change in peak area across the four timepoints (one-way ANOVA, FDR-corrected p-value ≤ 0.05, Benjamini-Hochberg, n = 4) and a FC (expressed in log2) relative to 0 h of either ≥ 3 (formed) or ≤ -1 (depleted) at one or more timepoints; (iii) a significant difference from the abiotic control, retaining features whose time course in the culture differed from the sterile control (two-way ANOVA on the species × timepoint interaction, FDR-corrected p-value ≤ 0.05, Benjamini-Hochberg, n = 4), thereby excluding abiotic degradation products and media background; (iv) removal of measurement artefacts, comprising low-intensity features (mean peak area < 500 counts at all timepoints) and in-source fragment ions, identified as co-eluting ions of lower *m/z* whose intensity profiles correlated with a higher-mass ion (Pearson r ≥ 0.7). This workflow yielded a total of 97 unique steroid metabolites across 125 detected biotransformation reactions, associated with 16 steroids and 11 bacterial species and 44 steroid-bacteria combinations (Supplementary Table S6). To infer the underlying chemical modifications, we calculated the mass difference between each putative metabolite and the original steroid compound, which indicates likely differences in atomic composition. On this basis we assigned distinct biotransformation reactions and their combinations to 38 of the 125 detected biotransformations (30%), distributed across 24 of the 44 steroid-bacteria pairs. Additionally, we predicted molecular formulas using SIRIUS (version 5.8.6) and assigned putative structures for 38 of the 97 unique steroid metabolites (COSMIC confidence score ≥ 80)^36, 37^ (Supplementary Table S7).

In the MS/MS acquisition we prioritized steroid-bacteria combinations with substantial depletion of the steroid drug (>50%). To examine whether the biotransformation reactions and metabolites identified in this primary dataset were shared across the bacterial panel, we performed an additional suspect-screening round, profiling this set of metabolites across steroid-bacteria combinations with lower drug conversion. This revealed several biotransformation reactions shared across multiple bacteria. For instance, side-chain cleavage to the corresponding 17-ketone was detected for both dexamethasone and flumethasone, in each case by both *C. scindens* and *S. herbicidovorans* (Fig. 2b, Supplementary Fig. 3a). Reduction of exemestane and the combined hydrolysis and dehydrogenation of fludrocortisone acetate were likewise observed across more than one species (Fig. 2c-d, Supplementary Fig. 3b-c). Through manual curation we expanded the dataset from 125 to 167 detected biotransformations and increased the proportion with an assigned reaction to 47% (79/167) of tested steroids. Altogether, this resulted in metabolite detections in 20 additional steroid-bacteria combinations and covering 18 steroids and 11 bacterial species (Fig. 2e-f). Leveraging the time-resolved experimental design, we classified each biotransformation product according to its abundance trajectory across the four timepoints. Of the 167 detected biotransformation reactions, 32 corresponded to initial products that occur at t0 and decline over time, 65 to transient intermediates that rose and subsequently decreased, and 70 to terminal products that accumulated (Supplementary Table S7). This kinetic classification enabled a more detailed reconstruction of sequential biotransformation steps, such as the stepwise deacetylation of fludrocortisone acetate and following oxidation to 9-fluoroprednisolone by *R. erythropolis* (Supplementary Fig. 3d).

Among the tested bacteria, *S. herbicidovorans* stood out for its broad capacity to transform steroids. It produced the highest number of metabolites in the screen (40 unique metabolites, compared with 37 for *C. testosteroni* and 32 for *R. erythropolis*), acting on 13 distinct steroid substrates. Like *C. scindens*, it catalyzes the desmolase-type side-chain cleavage, but unlike *C. scindens* it additionally transformed esterified prodrugs, and catalyzed several redox biotransformation reactions (Fig. 2e). Given this broad activity, we next set out to systematically characterize the steroid-transforming capacity of *S. herbicidovorans*, to define its substrate scope and to identify steroid-transforming enzymes.

### *S. herbicidovorans* displays diverse steroid biotransformation activity

Although *Sphingobium* species are known for versatile steroid biotransformation capacity, most of these biotransformation reactions are associated with bile acids degradation^38, 39^, while knowledge for other steroid modifications is limited. In our screen, *S. herbicidovorans* not only exhibited a desmolase-like activity but also generated the greatest diversity of biotransformation products, yielding 40 distinct metabolites derived from 13 steroid substrates (Fig. 2f). To evaluate substrate scope, we tested the activity of *S. herbicidovorans* with an additional panel of corticosteroids (eight additional compounds plus fludrocortisone acetate from the initial screen), selected to vary defined features of the cortisol scaffold, including the C1-C2 double bond, the C11 oxidation state, the C17-hydroxyl and its associated side chain (Supplementary Table S8). We performed the biotransformation assays as described above, with sampling at 0.5, 1, 2, 4, 8, 24 and 48 h. Fludrocortisone acetate was rapidly hydrolyzed to fludrocortisone within one hour, and fludrocortisone, along with the other tested compounds, was completely biotransformed within 24 h (Fig. 3a, Supplementary Fig. 4a). Building on the side-chain cleavage, reduction and oxidation activities identified in the initial screen, we applied suspect screening to the extended steroid panel, searching for the products of these reactions and their combinations. This identified 32 unique biotransformation metabolites (Supplementary Table S9). The time-resolved data resolved sequential steps within these pathways. For fludrocortisone, the denser time course resolved several downstream intermediate products of the side-chain cleavage that were not distinguished in the primary screen. Under the four-timepoint sampling, the product of hydrolysis, side-chain cleavage and oxidation had appeared to accumulate, whereas the higher resolution showed that, following hydrolysis, side-chain cleavage yields a transient product that undergoes further modification over time (Supplementary Fig. 4b). Additionally, for some substrates such as fludrocortisone and hydrocortisone, the same steroid gave rise to divergent products through parallel routes, in which side-chain cleavage and redox modification acted in different order or combination, resulting in more than one metabolic fate (Supplementary Fig. 4b-c). Across this panel, *S. herbicidovorans* transformed all tested corticosteroids, catalyzing side-chain cleavage and redox reactions. The biotransformation of these additional substrates further emphasizes the versatile biotransformation capacity of *S. herbicidovorans* and suggests the underlying enzymatic machinery as an attractive target for further mechanistic characterization and for applications in steroid biotechnology.

**Fig. 3.**
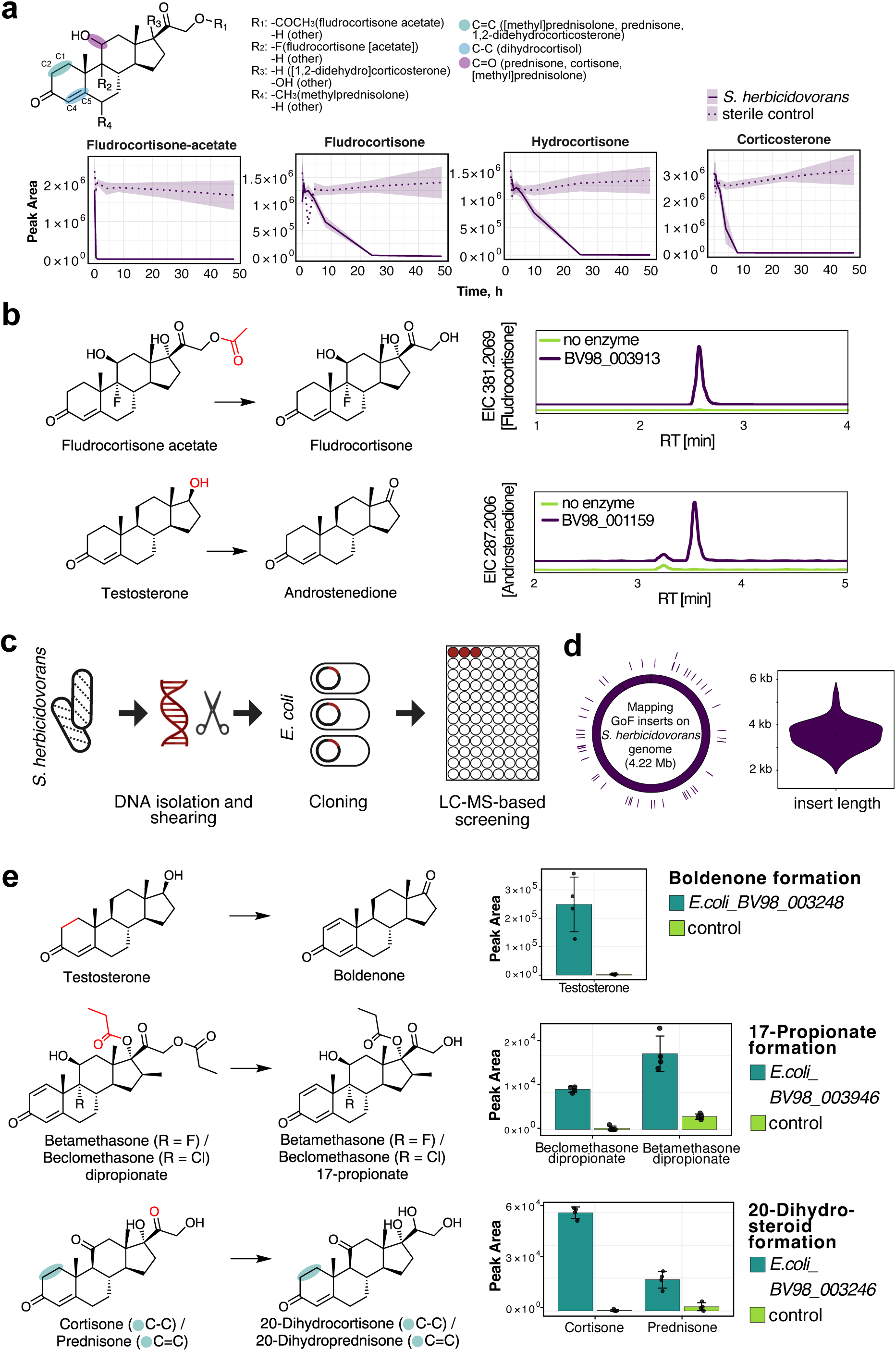
Steroid biotransformation in *S. herbicidovorans* and identification of involved genes. **a,** General structure of tested corticosteroids and degradation curves upon incubation with *S. herbicidovorans*. **b,** Biotransformation genes inferred from homology and product EICs: hydrolysis of fludrocortisone acetate upon incubation with purified hydrolase BV98_003913 and oxidation of 17-hydroxyl group of testosterone by purified oxidoreductase BV98_001159. **c,** Scheme of genetic gain-of-function library construction and screening pipeline. **d,** Mapping of DNA inserts onto the *S. herbicidovorans* genome, indicating genomic coverage and insert distribution. **e,** Examples of biotransformation identified with gain-of-function screening approach and product formation by *E. coli* expressing respective genes (n = 4 replicates, 16 h incubation; error bars, mean ± SD): hydrolysis of betamethasone dipropionate and beclomethasone dipropionate to respective 17-monopropionate by hydrolase *BV98_003946*, reduction of cortisone and prednisone side chain by 20-ketosteroid hydrogenase *BV98_003246*, 1,2-dehydrogenation of testosterone by dehydrogenase *BV98_003248*.

To identify enzymes responsible for steroid biotransformation in *S. herbicidovorans*, we first employed sequence homology searches, focusing on activities typically associated with aerobic steroid metabolism. Among aerobic organisms, diverse steroid-transforming enzymes including pregnane side-chain cleavage have been described in the fungal demethoxyviridin biosynthetic pathway^40^. We examined whether isofunctional enzymes might be present in *S. herbicidovorans*, and identified two proteins with a sequence identity higher than 25%: hydrolase *BV98_003913* (26% sequence identity with vidP) and oxidoreductase *BV98_001159* (25% sequence identity with vidH). We validated that *BV98_003913* catalyzes the hydrolysis of fludrocortisone acetate, and *BV98_001159* catalyzes the oxidation of the 17-hydroxyl group of testosterone to form the 17-keto androstenedione using purified enzymes produced with a cell-free expression system (Fig. 3b, Supplementary Fig. 5a-b).

Despite successful identification of enzymes using homology searches, it remains challenging due to the limited number of experimentally validated reference sequences. To further expand the list of steroid-transforming enzymes, we applied a gain-of-function (GoF) screen using a genetic library to identify DNA fragments of *S. herbicidovorans* that confer steroid biotransformation activity when expressed in *E. coli*. First, genomic DNA of *S. herbicidovorans* was extracted and sheared to 2–6 kb fragments, followed by cloning them into an *E. coli* expression vector and arrayed 30,000 transformed *E. coli* clones in a 384-well format (Fig. 3c). Sequencing of 48 randomly selected plasmid inserts revealed a mean insert length of 3.5 kb, which suggested an 18-fold genome coverage for the entire expression library (Fig. 3d). Second, we assembled 84 pools of 384 clones, incubated them with a mixture of the steroids metabolized by *S. herbicidovorans*, and measured steroid and metabolite levels over time to identify active library plates that included clones that gained specific steroid-metabolizing capacities. Third, we pooled rows and columns of active library plates, and repeated the steroid biotransformation metabolism assay to identify the plate position (bacterial clone) that gained a specific biotransformation activity, and then sequenced the genomic DNA inserts carried by these active clones. To validate the identified enzymes, we PCR-amplified respective gene sequences, cloned them in *E. coli* and tested the expression clone for steroid-metabolizing activity with selected compounds. Following this approach, we identified and validated three additional steroid-biotransformation enzymes: 1,2-dehydrogenase that converts testosterone into boldenone (*BV98_003248*), hydrolase of beclomethasone dipropionate and betamethasone dipropionate (*BV98_003946*), and 20-ketosteroid hydrogenase that reduces 20-keto group of cortisone and prednisone (*BV98_003246*) (Fig. 3e, Supplementary Fig. 5c-e). Thus, gain-of-function screens can systematically link genes to reactions and products independent of prior mechanistic annotation. However, our inability to identify the steroid side-chain cleavage enzyme despite combining genetic screens with targeted homology searches demonstrates the limitations of heterologous expression- and sequence homology-based approaches and the need for alternative strategies.

### Discovery of novel side-chain cleavage enzyme using proteomics

We observed steroid side-chain cleavage by *S. herbicidovorans* for fludrocortisone, hydrocortisone, cortisone, prednisone, methylprednisolone, which possess a hydroxyl group at the C17 position, as well as for corticosterone, which lacks it (Fig. 4a, Supplementary Fig. 6a). Previous studies have shown that bacterial steroid-metabolizing enzymes are frequently induced in the presence of their steroid substrates, enabling the identification of pathway-associated genes and enzymes through comparative expression analyses^24, 41^. We therefore tested whether exposure of *S. herbicidovorans* to synthetic steroids induced a distinct proteomic response that would highlight candidate steroid-transforming enzymes. To this aim, we cultured *S. herbicidovorans*, added either compounds - fludrocortisone acetate, fludrocortisone, corticosterone (Fig. 4c), cortisone and hydrocortisone (not shown) - or vehicle control and incubated them for one hour, followed by cell pelleting, protein extraction, and quantitative proteome analysis (Fig. 4b). We detected peptides mapping to 2,770 proteins of *S. herbicidovorans*. Comparative proteomic analysis revealed two proteins, BV98_003899 and BV98_001162, that were consistently more abundant in the presence of steroids relative to the vehicle control (log2 FC > 1, limma^42^ on vsn-normalized^43^ TMT intensities, Benjamini-Hochberg-adjusted p ≤ 0.01), suggesting an adaptive response linked to steroid transformation. Both proteins were significantly induced upon exposure to all five tested steroids. Gene *BV98_003899* encodes N5,N10-methylene tetrahydromethanopterin reductase, a distant homologue of LuxA luciferase from marine bacteria *Enhygromyxa salina*^44^ (29% sequence identity), and *BV98_001162* encodes a flavin-containing monooxygenase, which is homologous to a cyclohexanone monooxygenase^45, 46^ (37% sequence identity) and a steroid monooxygenase^47^ (33% sequence identity) from *Rhodococcus sp.* To test the activity of these proteins, we PCR-amplified the respective genes, cloned them into an *E. coli* expression vector and incubated the clone to test for steroid biotransformation with selected compounds. *E. coli* expressing BV98_003899 (hereafter *Sh*DES) exhibited activity toward 17-hydroxylated corticosteroids, producing the corresponding 17-keto metabolites. No side-chain cleavage products were detected with corticosterone or 1,2-didehydrocorticosterone, highlighting the importance of 17-hydroxyl group for binding and/or catalysis (Fig. 4d). Although it is likely that BV98_001162 is involved in the oxidation of the steroid side chain, we were unable to reconstitute its activity in *E. coli*.

**Fig. 4.**
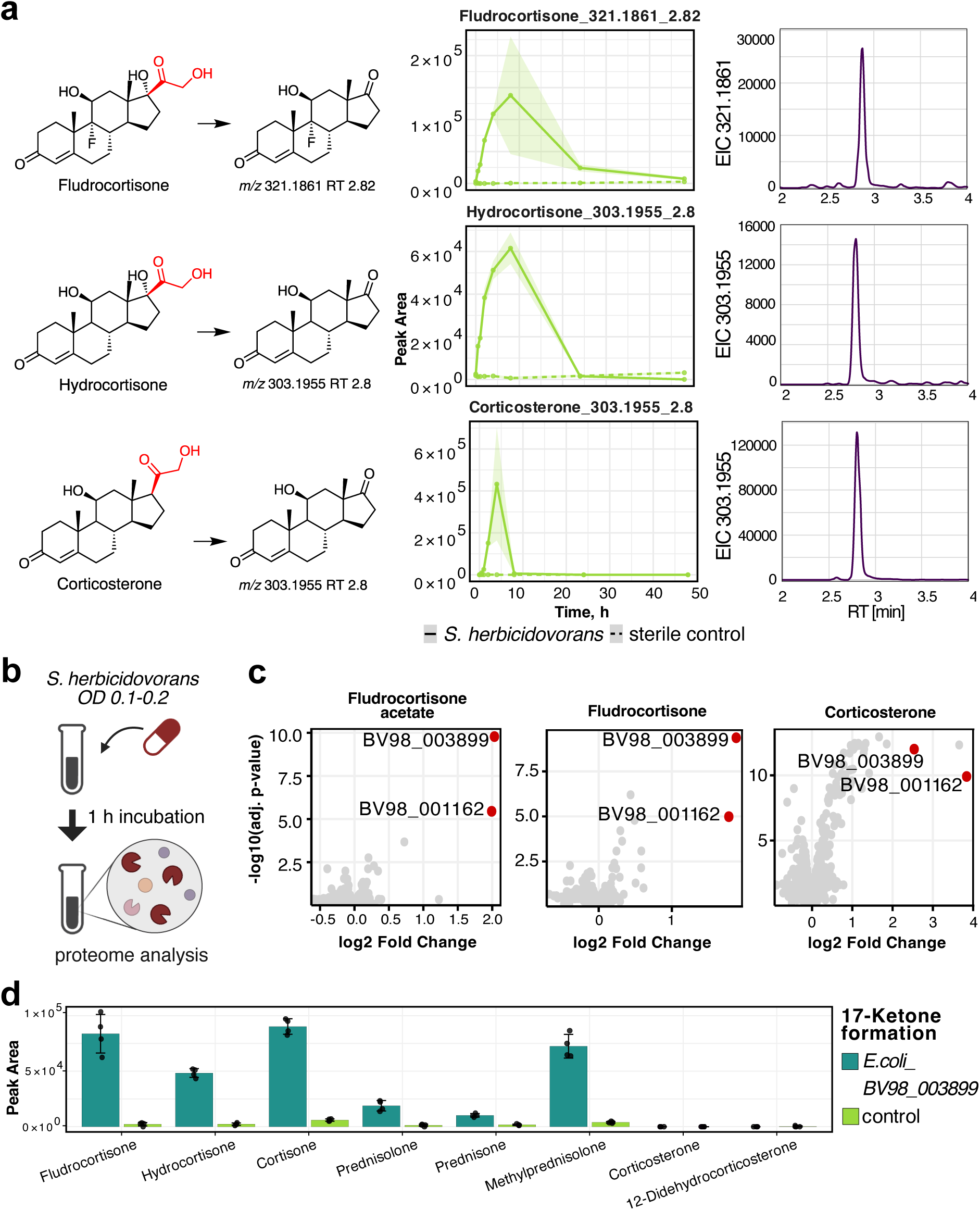
Desmolase reaction in *S. herbicidovorans*. **a,** Side-chain cleavage of fludrocortisone, hydrocortisone and corticosterone in *S. herbicidovorans* leads to the production of 17-ketone metabolites, shown as time-resolved formation and depletion of respective 17-ketone product (n = 4 replicates; shaded area, mean ± SD), alongside representative extracted-ion chromatograms (EICs). **b,** Schematic overview of the experimental proteomics workflow for gene identification. **c,** Analysis of expression proteomics highlighting differentially abundant proteins upon compound (fludrocortisone acetate, fludrocortisone and corticosterone) *vs* vehicle (DMSO) treatment (n = 3 replicates; limma on vsn-normalized TMT intensities, FDR-corrected p-value ≤ 0.01, Benjamini-Hochberg). **d,** Validation of *BV98_003899* gene activity by expression in *E. coli*: formation of side-chain cleavage metabolites (n = 4 replicates, 16 h incubation; error bars, mean ± SD).

The activity of *S. herbicidovorans* to cleave the corticosterone side chain fundamentally distinguishes it from *C. scindens*, which instead only reduces the C20-keto group to a 20-hydroxyl group^48^. In *S. herbicidovorans*, corticosterone biotransformation yields the side-chain cleavage product with the same retention time as observed in hydrocortisone metabolism (Fig. 4a). We propose that side-chain cleavage of corticosterone may proceed via an initial oxidative step, however, screening for the corresponding oxidation intermediate did not reveal formation of hydrocortisone. Instead, we detected a metabolite with the same exact mass as hydrocortisone but a distinct retention time, consistent with an isomeric oxidation product rather than canonical 17α-hydroxylation product (Supplementary Fig. 6b). Together, these findings suggest that *S. herbicidovorans* encodes a previously unrecognized corticosteroid side-chain cleavage pathway, including a novel desmolase enzyme in aerobic bacteria. Next, we examined whether steroid biotransformation activities like desmolase contribute to community-level steroid metabolism and metabolic cross-feeding across microbial communities in environmental contexts.

### Cross-feeding of steroid metabolites by gut and environmental bacteria

Our biotransformation screen using axenic bacterial cultures revealed that steroid compounds frequently undergo distinct biotransformation reactions by different bacterial species, resulting in diverse steroid metabolites. In nature, where bacteria rarely exist in isolation but rather in communities, products of these biotransformations can serve as substrates for subsequent reactions in neighboring bacteria, enabling community-level steroid metabolism through metabolic cross-feeding. For example, *C. scindens* produced metabolites of lytic side-chain cleavage with dexamethasone, 1,2-dihydro-dexamethasone, betamethasone, flumethasone, and triamcinolone. However, esterified corticosteroids (*i.e.,* fludrocortisone acetate and dexamethasone phosphate) are not biotransformed by *C. scindens*. Among gut-associated bacteria, however, hydrolysis of the latter is performed by *B. uniformis* and *P. vulgatus*, raising the possibility that these organisms enable downstream metabolism of de-esterified products and facilitate subsequent biotransformation, such as *C. scindens*. To test steroid biotransformation as a result of bacterial cross-feeding, we conducted subsequent cultivation experiments in which fludrocortisone acetate was first incubated with *B. uniformis*, followed by a second incubation of the sterile filtrate with *C. scindens* (Fig. 5a). As expected, we only detected the product of the desmolase reaction when the compound underwent sequential incubation with both species, while incubation of fludrocortisone acetate with only *B. uniformis* or only *C. scindens* led to either hydrolysis or intact fludrocortisone acetate, respectively (Fig. 5b). Similarly, we tested cross-feeding of dexamethasone phosphate with *P. vulgatus* and *C. scindens* (Supplementary Fig. 7a-b). Both *B. uniformis* and *P. vulgatus* are highly prevalent and abundant in the human gut, present in nearly every individual. Their prevalence suggests that hydrolysis of prodrugs such as fludrocortisone acetate is a common activity in the microbiota, whereas *C. scindens* occurs more rarely in fecal metagenomes.

**Fig. 5.**
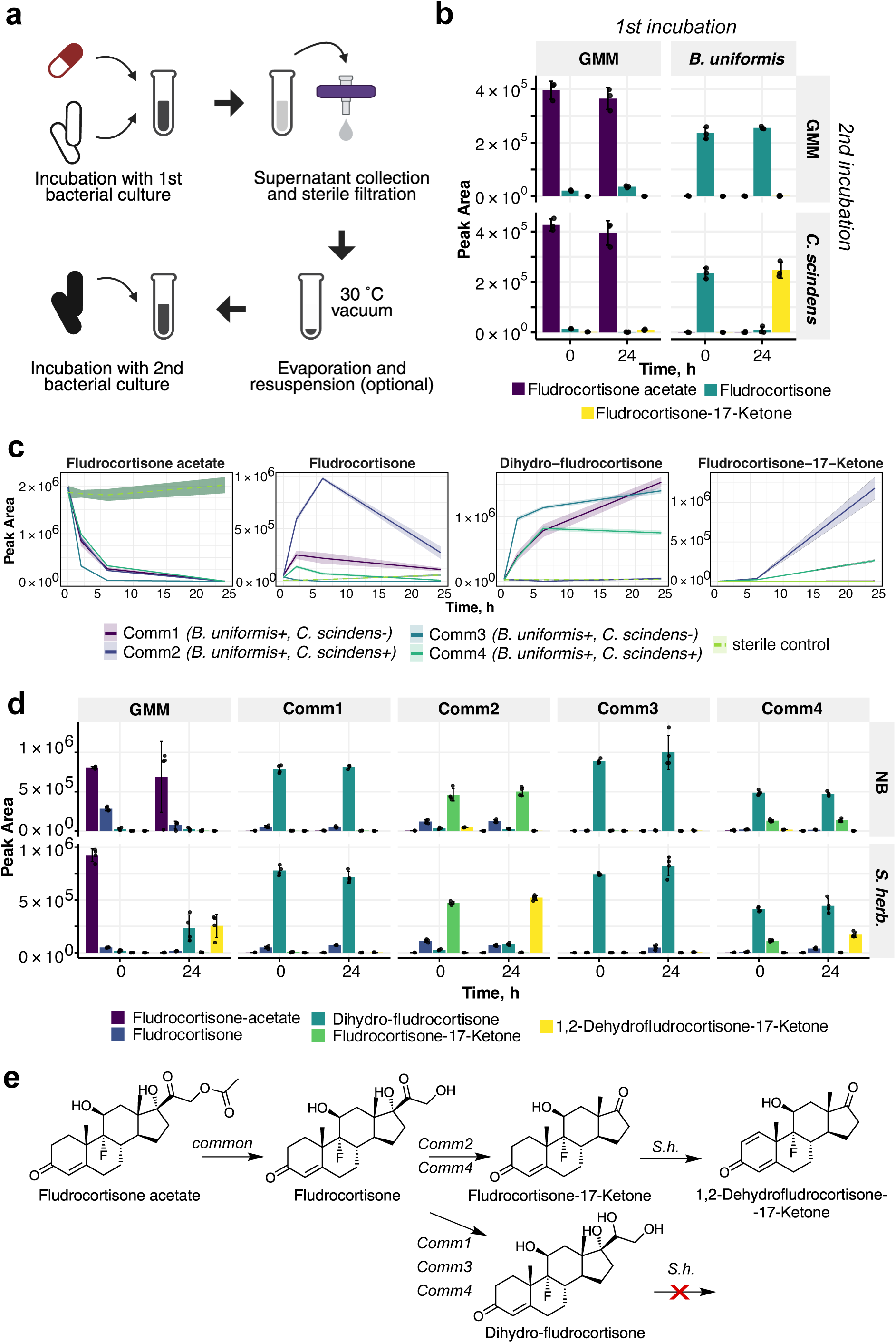
Cross-feeding of fludrocortisone acetate in single species and microbial communities. **a,** Schematic overview of the cross-feeding assay including first incubation, supernatant filtration and evaporation and second incubation. **b,** Cross-feeding of fludrocortisone acetate between gut bacteria *B. uniformis* and *C. scindens* (n = 3 replicates; first incubation 48 h; cross-feeding incubation 24 h; error bars, mean ± SD); GMM is a sterile media control. **c,** Fludrocortisone acetate conversion by four human-stool derived bacterial communities and formation of respective metabolites (n = 4 replicates per community; shaded area, mean ± SD). **d,** Cross-feeding of fludrocortisone acetate metabolites produced by gut bacterial communities and *S. herbicidovorans;* (n = 4 replicates, 24 h incubation; error bars, mean ± SD); GMM and NB are sterile media controls; **e,** Scheme of fludrocortisone acetate biotransformation by gut bacterial communities and *S. herbicidovorans*.

To demonstrate steroid biotransformation based on bacterial cross-feeding in human-stool derived bacterial communities, we next selected four fecal communities from healthy donors (Comm1-Comm4, Supplementary Table S11). All communities were subjected to metagenomic analysis in a previous study^49^: whereas all communities contained *B. uniformis, C. scindens* was detected only in Comm2 and Comm4. As expected, over 24 h incubation of fludrocortisone acetate with these communities the product of the desmolase reaction (fludrocortisone-17-ketone) was observed exclusively in Comm2 and Comm4, indicating that this metabolic step depends on the presence of specific community members. Another product (20-dihydrofludrocortisone) was observed in Comm1, Comm3 and Comm4, which is likely produced by 20-hydroxysteroid dehydrogenases reported for intestinal bacteria^50, 51^ (Fig. 5c). Based on similar cooperative metabolism, we have also shown that dexamethasone-17-ketone can be produced through hydrolysis by *P. vulgatus* and subsequent side-chain cleavage by *C. scindens* (Supplementary Fig. 7c). Overall, these findings demonstrate that terminal steroid biotransformation products are determined by community composition and metabolically cooperative interactions between distinct bacterial members.

To test whether cooperative steroid biotransformation extends beyond the gut to cross-ecosystem metabolism, we profiled MGnify metagenomes database (>2.4 billion non-redundant protein sequences across 297 different biomes) for homologues of the two desmolases: *Sh*DES from *S. herbicidovorans* and DesA from *C. scindens*. The two enzymes occupied distinct ecological niches. *Sh*DES-like proteins were broadly distributed across environmental habitats and nearly absent from gut-associated microbiomes, whereas DesA homologues were found in more variable ecosystems, including the human gut (Supplementary Fig. 8a-b). In fact, *Sh*DES homologues were present in only a subset of *Sphingobium* species (Supplementary Fig. 8c) and, across soil metagenome-assembled genomes, were largely confined to the *Pseudomonadota* and *Actinomycetota* (Supplementary Fig. 8d). These distributions define two ecologically distinct protein families, separating host-adapted anaerobic steroid metabolism from its aerobic environmental counterpart.

We next asked whether these two systems can functionally interface, using a desmolase complementation assay in which *S. herbicidovorans* was supplied with metabolites generated by gut communities. Gut community cultures were incubated with fludrocortisone acetate, and their supernatants were collected, sterile-filtered, evaporated to remove residual medium, resuspended in fresh medium and mixed with *S. herbicidovorans* (Fig. 5d). Side-chain cleavage products were detected in Comm2 and Comm4 supernatants after exposure to *S. herbicidovorans*. In these cases *S. herbicidovorans* drove the metabolite to an endpoint distinct from the one of *C. scindens*: alongside the side-chain cleavage product fludrocortisone-17-ketone (m/z 321.1861, RT 2.82; the *Sh*DES product), we detected an oxidized product at m/z 319.1704 (RT 2.72), consistent with A-ring oxidation accompanying side-chain cleavage, together with a 17-ketone isomer at m/z 321.1861 (RT 2.58) (Supplementary Fig. 4b, Supplementary Fig. 9a). The same endpoints arose when fludrocortisone acetate was cross-fed through *B. uniformis* and/or *C. scindens* before exposure to *S. herbicidovorans* (Supplementary Fig. 9b), confirming that they mark the characteristic endpoint of *S. herbicidovorans* rather than of the fecal community. By contrast, no desmolase products were detected with supernatants from Comm1 and Comm3, in which the predominant residual metabolite was 20-dihydrofludrocortisone (Fig. 5d-e), suggesting that reduction of the C20-keto group blocks subsequent desmolase activity. Together, these results show that steroid biotransformations within one microbial niche can generate metabolites that are further modified by bacteria in another ecosystem. At the same time, these transformations by microbiota from one ecosystem can reduce accessibility to downstream reactions by altering key structural features required for enzyme activity.

## DISCUSSION

We systematically mapped synthetic steroid biotransformation across gut and environmental bacteria, revealing a broad range of reactions spanning hydrolysis, redox transformations, and steroid side-chain cleavage. By combining homology-guided discovery, genetic gain-of-function screening, and quantitative proteomics, we identified six distinct steroid-transforming enzymes despite low sequence conservation and divergent metabolic functions. In particular, we identified a previously uncharacterized aerobic desmolase, extending beyond classical anaerobe *C. scindens* activity to aerobic environmental bacteria. We further demonstrated that these activities operate in natural bacterial communities, where cooperative metabolism and cross-feeding enable sequential steroid transformations across species. Collectively, these findings establish steroid metabolism as a property of microbial communities, in which complementary enzymatic activities across species drive stepwise and alternative transformation pathways. Such cooperative metabolism is likely to influence both host exposure and environmental persistence of synthetic corticosteroids, progestins, and estrogens, particularly for compounds resistant to abiotic degradation. Moreover, side-chain cleavage products generated from corticosteroids by gut- or environment-associated desmolase reaction may provide additional metabolic entry points that facilitate downstream degradation in activated sludge and decrease environmental persistence^13^. Introducing *S. herbicidovorans* or its desmolase into activated sludge could, in principle, supply this activity through bioaugmentation, though stable establishment and persistence of introduced strains at full scale remains a recognized challenge^52, 53^. Beyond the ecological relevance, side-chain cleavage of this type is also of long-standing industrial interest, as the microbial conversion of abundant and low-cost phytosterols into the 17-ketosteroids provides the key androstane precursors for the semisynthesis of most commercial steroid drugs^54^. Therefore, the identified enzymes expand the known catalytic repertoire for steroid scaffold remodeling and provide promising enzymatic tools for biocatalytic steroid functionalization and bioremediation of persistent steroid pollutants.

## MATERIALS AND METHODS

### Chemicals

Tested compounds were purchased from Sigma/Merck, Biomol, TCI or Steraloids as listed in Supplementary Tables S2 and S8. LC-MS-grade solvents were purchased from Guyer. Other chemicals were purchased from Sigma/Merck if not otherwise stated.

### Bacterial strains

List of bacterial strains used in this study are listed in Supplementary Table S1.

### Molecular cloning

Plasmids and primers used in the study are listed in Supplementary Table S10.

### Biotransformation screening

#### Gut bacteria

Biotransformation assays with gut bacteria were carried out anaerobically using an anaerobic chamber (Coy Laboratory Products) with a gas mix of 20% CO_2_, 10% H_2_, and 70% N_2_. Frozen glycerol stocks of bacterial strains were plated on brain-heart infusion (BHI; Becton Dickinson) agar with 10% horse blood and incubated at 37°C for ∼48 h, then a single colony was picked and inoculated into 10 mL of gut microbiota medium (GMM)^55^, and grown at 37°C for ∼16 h. Bacterial cultures were then diluted 1:10 (v/v) with 50% GMM medium (GMM: sterile water 1:1), added onto 96-well plate containing the tested drugs in 4 replicates at a final concentration of 5 μM in a volume of 160 μL, and incubated anaerobically at 37°C. Samples (20 μL) were collected at 0, 3, 9 and 27 h and stored at -80 °C until further processing for analysis by LC-MS (see ‘Mass spectrometry analysis’ section).

#### Environmental bacteri

Biotransformation assay with environmental bacteria was carried out aerobically. Frozen glycerol stocks of bacterial strains were plated on lysogeny broth (LB) agar plates at 30°C for ∼48 h, then a single colony was picked and inoculated into 10 mL of DSMZ 1 medium (Nutrient Broth, NB) supplemented with ATCC Trace Minerals and ATCC Vitamin Supplement mix, and grown at 30°C for ∼48 h. Bacterial cultures were then diluted 1:10 (v/v) with 20% NB medium (NB: sterile water 1:4), added onto 96-well plate containing the tested drugs in 4 replicates at final concentration of 5 μM in a volume of 160 μL, and incubated aerobically at 30°C and shaking at 180 rpm. Samples (20 μL) were collected at 0, 3, 9 and 27 h and stored at -80 °C until further processing for analysis by LC-MS (see ‘Mass spectrometry analysis’ section). For testing additional steroid compounds with *S. herbicidovorans,* the assay was carried out the same way, apart from the final volume of the assay was 200 μL, and samples (20 μL) were collected at the timepoints 0, 0.5, 1, 2, 4, 8 and 24 h.

### Mass spectrometry analysis

#### Metabolite extraction

Liquid samples were processed for LC-MS analysis using an organic solvent extraction method, following a protocol similar to one previously described^22^. In brief, a mixture of acetonitrile:methanol (1:1) was added to the samples (5:1 v/v), followed by the addition of an internal standard mix, containing sulfamethoxazole, doxycycline, nafcillin, and diclofenac, each at a final concentration of 0.4 μM. The mixtures were then incubated at -20°C for at least 1h and centrifuged at 4300 rcf for 15 min at 4°C. The supernatant was then collected, diluted with water (1:1) and injected into the LC-MS system.

#### LC-MS analysis and MS1 acquisition

Analysis was performed using reversed-phase chromatography with an InfinityLab Poroshell HPH-C18 column (2.1x100 mm, 1.9 μm) with an Agilent 6546 LC/qTOF system. Water with 0.1% formic acid and acetonitrile with 0.1% formic acid were used as mobile phases A and B, respectively, with a gradient 5 to 95% phase B in 5.4 min and a flow rate of 0.4 ml/min. The qTOF instrument was operated in positive scanning mode (100-1700 *m/z*). Online mass calibration was performed using a second ionization source with a continuous infusion (5 μL/min) of Agilent reference mass solution containing protonated purine (*m/z* 121.0509) and protonated hexakis(1H,1H,3H-tetrafluoropropoxy)phosphazine (HP-921, *m/z* 922.0098).

#### MS2 acquisition

LC-MS/MS was performed using the chromatographic separation and source parameters described above, and the auto-MS/MS mode of the instrument with a preferred inclusion list for ions of interest with *m/z* tolerance of 20 ppm, RT tolerance of 0.4 min, narrow isolation width (∼1.3 *m/z*) and collision energy of 10eV.

### Analysis of mass-spectrometry data

#### Targeted feature detection (MS1)

The MassHunter Quantitative Analysis Software (Agilent, version 10.0) was used for peak detection and integration based and *m/z* and RT values of chemical standards. Left and right *m/z* tolerance were set to 20 ppm, left RT tolerance was set to 0.2 min and right RT tolerance of 0.3 min. Peak integrations were manually curated.

#### Analysis of MS1 targeted data

The decrease of the extracted peak area was used to assess whether the steroid was biotransformed or not. To account for the fast metabolism, which was observed in several samples with *C. testosteroni* where the parent steroid compound was not detected at 0 h timepoint, peak area of the compound in the control (pH7) at 0 h was imputed instead. A two-way analysis of variance (ANOVA) test was performed to examine the interaction between peak areas across four timepoints and the control (pH7). The p-values were adjusted for multiple comparisons false discovery rate (FDR) using the Benjamini-Hochberg procedure (p.adjust function in R with method parameter set to “fdr”). To assess the biotransformation of the drug, percent of decreased peak area was calculated and filtered for significance (adjusted p-value ≤ 0.05).

#### Untargeted feature detection (MS1)

Untargeted analysis was carried out for selected steroid-bacteria pairs with parent compound depletion at one or more timepoints ≥50% together with 6 combinations below this threshold that were retained to probe for low-level metabolite formation (Supplementary Table S5). To this aim, mass spectrometry data were converted from the proprietary format (.d) to the *m/z* extensible markup language format (.mzML) using the command-line-based peak picking algorithm of ProteoWizard (version 3.0.23304)^56^ under WINE (version 7.8). The mzML files were then processed with MZmine version 3.3.0^57^. In short, feature detection and deconvolution were performed in a batch mode including following steps: mass detection, ADAP chromatogram builder (*m/z* tolerance 0.002\**m/z* or 10 ppm), local minimum resolver algorithm, 13C isotope filter, isotopic peaks finder, join aligner (*m/z* tolerance 0.002\**m/z* or 10 ppm and RT tolerance 0.1 min), feature list rows filter (minimum features in a raw 3, or feature present in 75% of replicates), peak finder and duplicate peak filter (full list of paraments is available in the batch file at https://github.com/m-beliaeva/Synthetic-steroids-metabolism-by-environmental-and-gut-bacteria). The results tables were extracted in Metaboanalyst-compatible .csv format.

#### Analysis of MS1 untargeted data and creation of MS/MS inclusion lists

Differential analysis was done using Metaboanalyst (www.metaboanalyst.ca) for each steroid-bacteria interaction. MZmine output file was loaded into ’one factor statistical analysis’ module and quantile normalization was applied. Differential analysis (DA) was performed in following pairs: 27 h vs 0 h, 9 h vs 0 h, 3 h vs 0 h. Pairwise comparisons were assessed using a t-test with Benjamini-Hochberg FDR correction for multiple testing. Thresholds were set to ≥ 1 for the fold change (FC, expressed in log2) and to ≤ 0.05 for the adjusted p-value. Metaboanalyst output files for each steroid-bacteria interaction were exported in .csv format, and the downstream processing was performed in R to finalize the lists of putative metabolites for MS/MS analysis. To this end, files from the DA for each steroid-bacteria interaction were merged and filtered to exclude metabolites present in DMSO control. Resulting data was then formatted to create include lists for MS/MS analysis. If the ion for the original compound was not present in the resulting inclusion list, it was added manually during the measurement.

#### MS2 library creation and spectral similarit

MS/MS data files were converted from the proprietary format (.d) to the *m/z* extensible markup language format (.mzML) using the command-line-based peak picking algorithm of ProteoWizard (version 3.0.23304)^56^ under WINE (version 7.8). The R package MergeION was used to create MS/MS libraries and calculate spectral similarity^35^. Individual MS/MS libraries were generated with the library generator function, applying preprocessing parameters including intensity normalization, a baseline of 200, and a maximum of 200 peaks. Spectral matching against steroid drugs and known hydrolytic metabolites (Supplementary Table S4) was performed using the library_query function with cosine and F1 similarity algorithms, requiring a minimum of 3 fragment matches and positive ion polarity, while allowing *m/z* and RT tolerances (mz_search = 0.002, ppm_search = 10, rt_search = 6). Consensus spectra were generated with a 0.005 *m/z* window to improve comparison accuracy. MS/MS spectra were saved in mascot generic format (.mgf) for the downstream analysis.

#### Filtering putative metabolites

For the putative hits with spectral similarity thresholds of 0.5 (cosine) and 0.1 (F1), the peak area was extracted from previously obtained untargeted data based on *m/z* and RT values. An ANOVA test was performed to examine the significance of area changes across four timepoints, and p-values were FDR-corrected (Benjamini-Hochberg). FC were then calculated for the time point pairs 27 h vs 0 h, 9 h vs 0 h, and 3 h vs 0 h. Molecular features with FC thresholds (log2 scale) of ≥ 3 (formed) or ≤ −1 (depleted) and adjusted p-value ≤ 0.05 were retained. The retrieved list of putative metabolites then underwent a two-way ANOVA test to examine the significant changes between peak areas across four timepoints and the control (pH7) in order to eliminate media components. Again, p-values were FDR-corrected (Benjamini-Hochberg), and filtered by adjusted p-value ≤ 0.05. All features with peak area ≤ 500 counts across all time points were excluded. To account for in-source ion-fragmentation, features with the same RT were tested for area pattern correlation (cor.test function in R returning Pearson’s correlation coefficient); if the correlation coefficient was ≥ 0.7, the feature with the smaller *m/z* value was excluded. Notably, in-source defluorination was a common artefact for fluorinated steroids, and respective entries were removed from the metabolite list. To suggest chemical formula and putative structure of the metabolites, SIRIUS software^36^ (version 5.8.6) and acquired MS/MS spectra in mgf format were used. The default parameters were used unless differently stated: for selected compounds the number of chorine, fluorine and nitrogen atoms were changed from the default value for a number of compounds based on the steroid molecule atomic composition for improved SIRIUS molecular formula prediction (Supplementary Table S7). CSI:FingerID^58^ was used for matching MS/MS spectra to databases, and structure match was ranked by COSMIC confidence score^37^.

### Cheminformatics

#### Chemical similarity

Chemical similarity analysis was performed using ChemmineR R package^59^. Compound structures were provided as SMILES codes, and functions smiles2sdf, sdf2ap and cmp.cluster were used to obtain pairwise distances of structural similarity.

#### Atomic composition calculations

Steroid formulas were calculated from SMILES codes using function MolFormFromSmiles.rcdk from RChemMass package^60^. Atomic differences between steroid drug and metabolite compounds were calculated using generate.formula function from rcdk R package^61^.

### Genome sequencing of *S. herbicidovorans*

#### Bacterial culture and DNA isolation

*S. herbicidovorans* was grown overnight in 5 mL of NB medium supplemented with ATCC Trace Minerals and ATCC Vitamin Supplement mix at 30°C for ∼48 h at 180 rpm. Bacterial cells were harvested by centrifugation, and genomic DNA was isolated using the ZymoBIOMICS DNA Miniprep Kit (D4300) according to the manufacturer’s instructions. Samples were bead-beaten in a TissueLyser II for 2 × 2.5 min at 30 Hz, with a 1 min break between runs. DNA was eluted in 100 µL of water pre-heated to 65 °C and incubated on the column for 10 min prior to centrifugation. DNA fragment length (∼26-30kb) was assessed using a FEMTO Pulse system. Samples were then submitted for PacBio HiFi sequencing at the EMBL Genecore in-house sequencing facility. *Quality control and assembly.* HiFi reads in FASTQ format were quality-assessed using SeqKit v2.10.0. Reads were filtered using a minimum quality threshold of Q20 and a minimum read length of 1 bp. Filtered HiFi reads were assembled using Flye v2.9.6 with plasmid and metagenomic settings enabled. Assembly completeness and contamination were evaluated using CheckM v1.2.2 under lineage-specific workflows.

#### Taxonomic assignment

Assembled contigs were separated into individual FASTA files using SeqKit. Taxonomic classification was performed using GTDB-Tk v2.4.0 with the corresponding Genome Taxonomy Database release. Unclassified contigs were retained for further evaluation as potential plasmids.

#### Plasmid identification

Putative plasmid sequences were identified using combination of two approaches. Mobile genetic elements, including plasmids and viral sequences, were identified using geNomad v1.11.1. In parallel, similarity screening against the PLSDB 2024_05_31_v2 plasmid database was conducted using Mash 2.3. Results from both approaches were integrated to flag candidate plasmid contigs.

#### Genome annotation

Gene prediction and functional annotation were performed using Prokka v1.14.6.

### *Sh*DES homologues mapping

#### MGnify mapping

Protein homologues were identified using the MGnify phmmer sequence search tool^62^ with the reference proteins *Sh*DES (UniProtID A0A086P4J8) and DesA (UniProt ID B0NC66). Only hits with E-values <1×10⁻⁶ were retained for downstream analysis (48,042 hits for *Sh*DES and 117,002 hits for DesA). The resulting MGYP protein identifiers were extracted and mapped against MGnify biome annotation tables. For each homologue, the associated biome categories and observation counts were retrieved from MGnify biome files. Biomes were grouped into broader ecological categories including gut-associated host, other host-associated, aquatic environment, terrestrial environment, waste or polluted environment, engineered or laboratory environments, and other or unknown environments. Relative prevalence was defined as the percentage of homologue observations belonging to each biome group within a given protein dataset. Normalized enrichment was calculated as the number of homologue observations in a biome group divided by the total number of MGnify observations associated with that biome group.

#### MGnify mapping across taxonomy-resolved soil protein catalogue

The MGnify soil protein catalogue v1.0, clustered at 100% amino acid identity, was downloaded from the MGnify genomes FTP repository (https://ftp.ebi.ac.uk/pub/databases/metagenomics/mgnify_genomes/). The catalogue contains 75,154,756 protein sequences predicted from 20,908 genomes, representing 19,472 prokaryotic reference species. Protein catalogue dataset was used as the reference database to identify putative *Sh*DES homologues using command-line NCBI BLASTP+ v2.17.0^63^. Hits were retained using the following criteria: ≥50% amino acid sequence identity, ≥50% query coverage, and an E-value <1×10⁻⁶. The *Sphingobium* core genome phylogeny and the precomputed bacterial phylogenetic tree provided with the MGnify soil genomes catalogue were midpoint-rooted and visualized using iTOL v7.2.1^64^.

#### Mapping across Sphingobium species

Complete genome assemblies of genus *Sphingobium* were retrieved from the NCBI Genome database using the NCBI command-line datasets tool v18.2.0 on 26 March 2026. In total, 63 complete RefSeq genomes were included in the analysis. All genomes were annotated using Bakta v1.12.0 with default parameters, resulting in the prediction of 284,503 protein-coding sequences in a custom database. A core genome alignment was generated from the 63 annotated genome files using Panaroo v1.5.2^65^. A maximum-likelihood phylogenetic tree was constructed using IQ-TREE v3.0.1 using the best-fit substitution model according to the Bayesian Information Criterion was GTR+F+I+R5 with 1,000 ultrafast bootstrap replicates. Putative homologues of the *Sh*DES were identified by querying the custom *Sphingobium* protein database using NCBI BLASTP+ v2.17.0.

### Cell-free protein synthesis and enzyme assays

#### Cell-free protein synthesis

Recombinant enzymes BV98_003913 and BV98_001159 were produced using a cell-free protein expression system (Nuclera). Gene constructs were assembled using eGene™ Prep Kit NC3008 and protein expression and purification was done using eProtein Discovery protocol. *Enzyme assay.* Enzymatic activity was evaluated in 50 µL total reaction volume in phosphate buffer saline (PBS, pH 7.4). To test hydrolase activity, 0.1 mM fludrocortisone acetate was incubated with 20 µM MgCl₂ and 2 µM enzyme BV98_003913. To test oxidoreductase activity 0.1 mM testosterone was incubated with 0.2 mM NAD⁺/0.2 mM NADP⁺ and 2 µM enzyme BV98_001159. All reactions were incubated for 16 h, 250 rpm. Reactions were terminated by quenching by 1:1 volume of ACN/MeOH mixture.

### Gain-of-function library screening of *S. herbicidovorans*

#### Preparing gain-of-function library

The preparation of heterologous expression library of *S. herbicidovorans* was adapted from^22^. Genomic DNA was extracted from ∼72 h culture of *S. herbicidovorans*. DNA was fragmented into 2-6 kb pieces using focused ultrasonication (Covaris E220 with miniTUBE red). DNA Fragments were subsequently cloned into a PCR-linearized pZE21 expression vector (primers 1 and 2, Supplementary Table S10) through blunt-ended ligation (Thermo Fisher Rapid DNA Ligation kit). Ligated products were run on a 0.5% agarose gel to isolate fragments between 5 and 10 kb, which were then purified with a gel extraction kit (Qiagen). Ligation products were transformed into *E. cloni* 10G Elite competent cells (Lucigen) via electroporation. Colonies that grew overnight were selected and arrayed in a 384-well plate containing LB medium with kanamycin, assisted by a colony-picking robot (Singer Rotor). Inoculated plates were incubated overnight at 37°C, then duplicated onto LB agar plates with kanamycin and stored at -70°C for preservation.

#### Gain-of-function screen

Screening of the gain-of-function library was carried out as previously described^22^. First all 384 colonies from a single library plate were pooled together and resuspended in 20% NB medium containing compounds of interest (final concentration 5 μM). The suspension was incubated aerobically at 37°C and shaking at 180 rpm. Samples (20 µL) were collected at 0, 0.5, 1, 2, 4, 8, 24, 48, and 72 h of incubation. Pools showing the ability to metabolize parent compound and/or to produce previously identified metabolite were then transferred to a new 384-well plate and cultured aerobically for 12 h at 37°C. Active pools were subsequently deconvoluted by re-pooling clones from the original 384-well plate by row and column using a liquid-handling platform (Agilent Bravo), followed by secondary screening of the resulting pools. Active clones were identified based on intersecting activity across row and column pools. Four individual colonies from active clones were isolated, re-evaluated for their metabolizing capability, and underwent Sanger sequencing to identify the genetic inserts using primers 3 and 4 (Supplementary Table S10).

### Expression proteomics of *S. herbicidovorans*

Pre-cultures of *S. herbicidovorans* were initiated from a single colony in NB medium supplemented with ATCC Trace Minerals and ATCC Vitamin Supplement and grown for 48 h at 30 °C. Cultures were then inoculated at an optical density (OD₆₀₀) of 0.1-0.2 into 60 mL of each medium and incubated overnight at 30 °C with shaking at 180 rpm. At mid-exponential phase (OD₆₀₀ ≈ 0.5), cultures were aliquoted into 2 mL tubes for proteomic analyses. Each tube was treated with 5 µL of 10 mM compounds or DMSO control, resulting in a final concentration of 25 µM. All conditions were prepared in triplicates. Cultures were incubated at 30 °C and 220 rpm for 60 min, harvested by centrifugation at 4 °C for 15 min, washed with 1 mL cold PBS, centrifuged again, and stored at −20 °C. Cell pellets were resuspended in 2% SDS in PBS and lysed by incubation at 98 °C for 10 min. Protein concentrations were measured using the BCA assay and 5 μg of protein per sample was digested following a modified SP3 protocol, as described previously^66, 67^. Eluted peptides were labeled with TMT18plex^68^ and were pre-fractionated under high pH conditions. Samples were then analyzed with liquid chromatography coupled to tandem mass spectrometry, as previously described^67^.

Raw files were converted to .mzML format using MSConvert from ProteoWizard, using peak picking, 64-bit encoding and zlib compression, and filtering for the 1000 most intense peaks. Files were then searched using MSFragger (v4.0)^69^ in FragPipe (21.1) against FASTA database uniprotkb_proteome_UP000024284_2024_01_12 containing common contaminants and reversed sequences. Data analysis was performed using R. Only proteins with at least one unique identified peptide were kept for the analysis. Protein intensities across all channels of each TMT set were normalized using variance stabilization normalization (vsn^43^) and differential abundance analysis was performed using linear mixed model analysis (limma^42^).

### Hit validation by targeted gene expression in *E. coli*

To validate identified steroid-metabolizing enzymes, gene sequences were PCR-amplified (primers in Supplementary Table S10), cloned into the pZE21 expression plasmid by Gibson cloning (NEBuilder HiFi DNA Assembly Kit, NEB), and electro-transformed *E. cloni* 10G Elite cells. *E. cloni* clones were inoculated into 5 mL of LB medium and incubated at 37°C, 250 rpm for 3 h aerobically, after which cultures were transferred to assay plates containing test compounds at a final concentration of 5 µM in 200 µL volume. Cultures were incubated aerobically at 37°C, 250 rpm and samples (20 µL) were collected at 0 h and 16 h. Samples were stored at -80 °C until further processing for analysis by LC-MS (see ‘Mass spectrometry analysis’ section).

### Cross-feeding assays

#### Cross-feeding assay with single species

Single colonies of *B. uniformis* and *P. vulgatus* were inoculated into 5 mL GMM (in three replicates) and grown anaerobically for ∼16 h. Cultures were subsequently diluted 1:1 (v/v) with sterile water, and compounds were added to a final concentration of 10 µM. Cultures were incubated anaerobically at 37°C, and samples were collected at 0 h and 48 h. Following incubation, cultures were filtered using 0.2 µm PVDF filter to remove bacterial cells. Collected supernatants were mixed 1:1 (v/v) with preculture of *C. scindens* (16 h preculture in GMM), followed by incubation and sampling at 0 h and 24 h. After 24 h samples were clarified by filtration. 100 µL of filtrate were dried under vacuum, and further used for the cross-feeding*. S. herbicidovorans* was inoculated into 10 mL of NB and grown at 30°C for ∼72 h at 180 rpm. Bacterial cultures were then diluted 1:1 (v/v) with NB medium, and added onto the plate with remaining dry residue to a final volume of 100 μL. Cultures were incubated aerobically at 30°C and shaking at 250 rpm. Samples were collected at 0 h and 24 h and stored at -80 °C until further processing for analysis by LC-MS (see ‘Mass spectrometry analysis’ section).

#### Cross-feeding assay with gut communities and S. herbicidovorans

Information about original gut communities, sample processing and metadata can be found in the original study^49^ (Supplementary Table 11). Anaerobic precultures were initiated inside an anaerobic chamber by inoculating 5 mL GMM with 1 mL of frozen community sample. Overnight cultures were diluted 1:1 (v/v) with GMM medium, added onto 96-well plate containing the tested drugs in 4 replicates at final concentration of 10 μM in a final volume of 1 mL, and cultures were incubated anaerobically at 37°C. Samples (20 µL) were collected at 0 h, 2 h, 6 h and 24 h. Cultures were then centrifuged, and supernatants were transferred to filter plates (Agilent 203982-100, 96-well plate with 0.2 µm PVDF filter) and clarified by filtration. 100 µL of filtrate were then dried under vacuum, and further used for the cross-feeding with *S. herbicidovorans. S. herbicidovorans* was inoculated into 10 mL of NB and grown at 30°C for ∼72 h at 180 rpm. Bacterial culture was then diluted 1:1 (v/v) with NB medium, and added onto the plate with remaining dry residue to a final volume of 100 μL. Cultures were incubated aerobically at 30°C and shaking at 250 rpm. Samples (20 µL) were collected at 0 h and 24 h and stored at -80 °C until further processing for analysis by LC-MS (see ‘Mass spectrometry analysis’ section).

## AUTHOR CONTRIBUTIONS

Conceptualisation and study design were carried out by M.A.B. and M.Z. Experimental investigation was performed by M.A.B., with gain-of-function and cloning work conducted by M.W.G. Proteomics analyses were performed by D.P. under the supervision of M.S. Whole-genome sequencing of *S. herbicidovorans* was carried out by A.S. Homology mapping was done by M.A.B. and A.S. The original draft of the manuscript was written by M.A.B., reviewed and edited by M.Z. All authors read the manuscript and contributed to critical revision.

## ACKNOWLEDGEMENTS

We acknowledge funding from EMBL. M.A.B. was supported by the EMBL Interdisciplinary Postdoctoral Programme under the Marie Skłodowska-Curie Actions COFUND, as well as by a Joachim Herz Add-on Fellowship. M.Z. acknowledges support from a European Research Council (ERC) Starting Grant (GutTransForm ID 10107835). We thank Jeanne Marechal for the help with biotransformation screening of gut bacteria, and we thank Kim Remans, Karine Lapouge and Yexin Xie at PepCore facility at EMBL for assistance with cell-free protein expression and purification.

## DATA ACCESS

The data are available in the main text or the supplementary materials. Genome sequencing and annotation of *S. herbicidovorans* are available in the European Nucleotide Archive (ENA) under study accession number PRJEB114257. The pre-processed metabolomics data are available on Zenodo (DOI 10.5281/zenodo.20523169) and R scripts for data analysis and figure generation are available on GitHub (https://github.com/m-beliaeva/Synthetic-steroids-metabolism-by-environmental-and-gut-bacteria).

---

**Supplementary Fig. 1.**
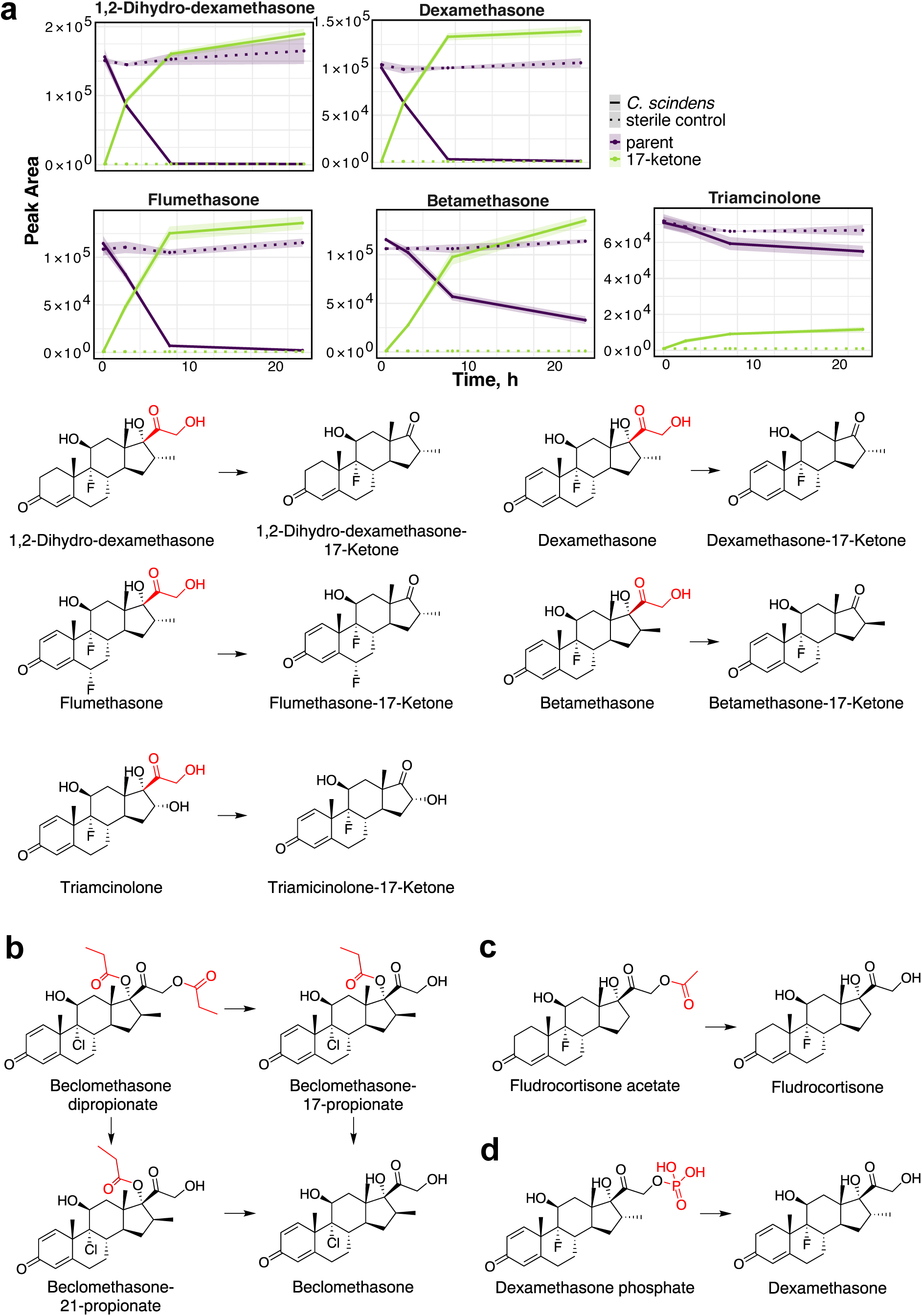
Detected metabolites of bacterial steroid biotransformation. **a,** Conversion of 1,2-dihydro-dexamethasone, dexamethasone, betamethasone, flumethasone and triamcinolone into respective 17-ketone derivatives by *C. scindens* (n = 4 replicates; shaded area, mean ± SD). **b,** Two-step beclomethasone dipropionate hydrolysis by *S. herbicidovorans*. **c,** Hydrolysis of fludrocortisone acetate by *B. uniformis*. **d,** Hydrolysis of dexamethasone phosphate by *P. vulgatus*.

**Supplementary Fig. 2.**
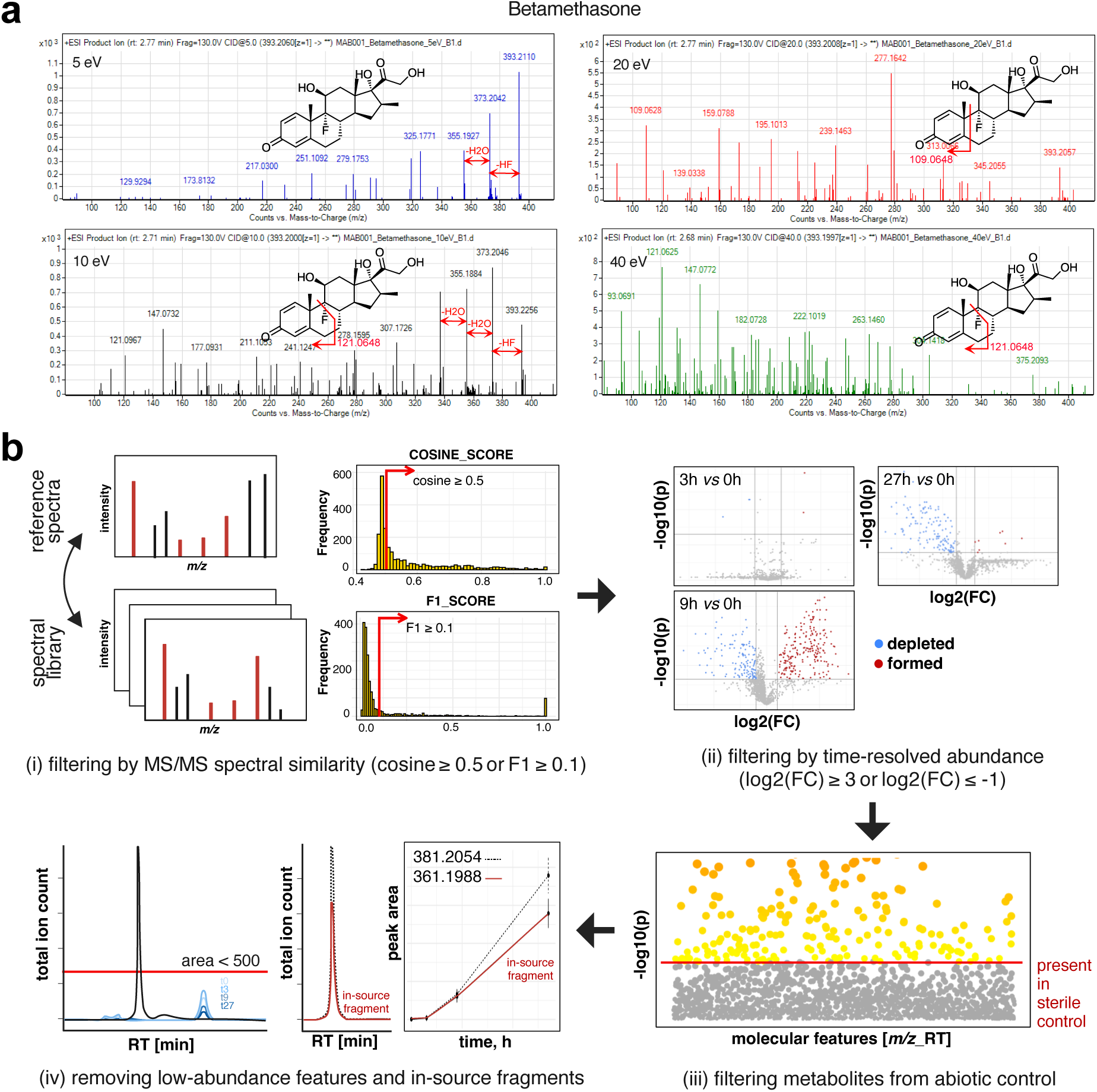
MS/MS fragmentation optimization and downstream metabolites analysis. **a,** Collision energy optimization (betamethasone example): at 5 eV the parent ion remains dominant and defluorination and dehydration fragments emerge; at 10 eV defluorination and dehydration products are observed together with lower *m/z* diagnostic fragments for the steroid core; at 20 and 40 eV increased fragmentation leads to progressive loss of larger structural units. **b,** Computational workflow for filtering putative biotransformation metabolites, applied in four sequential steps: (i) filtering by MS/MS spectral similarity to reference spectra (cosine ≥ 0.5 or F1 ≥ 0.1); (ii) filtering by time-resolved abundance at 3, 9 and 27 h relative to 0 h, retaining formed (log2(FC) ≥ 3) and depleted (log2(FC) ≤ −1) features; (iii) removal of features also present in the abiotic (sterile) control; and (iv) removal of low-abundance features (peak area < 500) and in-source fragments.

**Supplementary Fig. 3.**
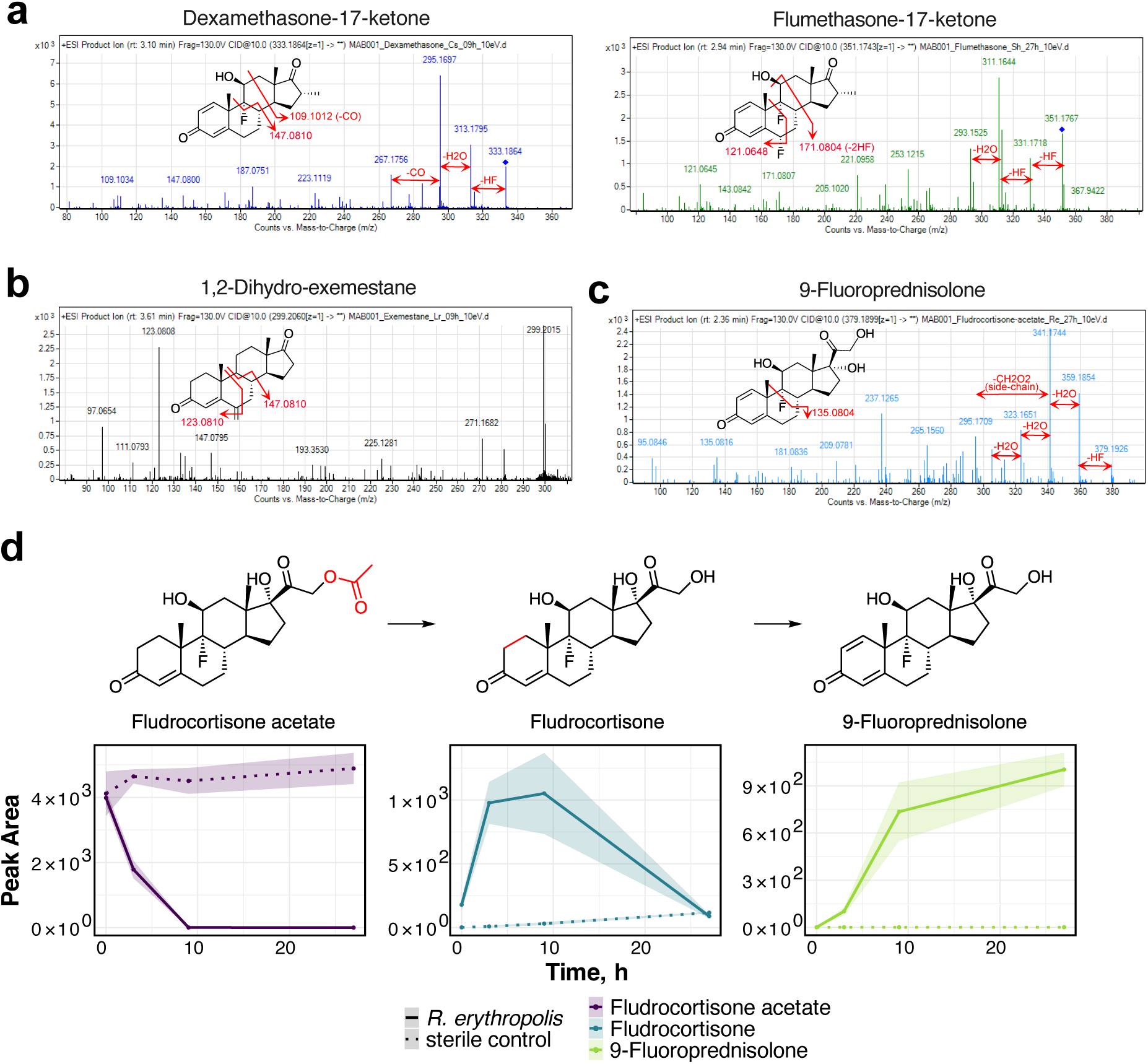
Products of bacterial steroid biotransformation and their molecular structure elucidation by MS/MS. **a-c,** Examples of MS/MS fragmentation of the dexamethasone-17-ketone and flumethasone-17-ketone produced by *C. scindens* and *S. herbicidovorans* (a), 1,2-dihydroexemestane produced by *L. reuteri* (b), and 9-fluoroprednisolone produced by *R. erythropolis* (c). **d,** Two-step biotransformation of fludrocortisone acetate by *R. erythropolis* including deacetylation to fludrocortisone and following oxidation to 9-fluoroprednisolone.

**Supplementary Fig. 4.**
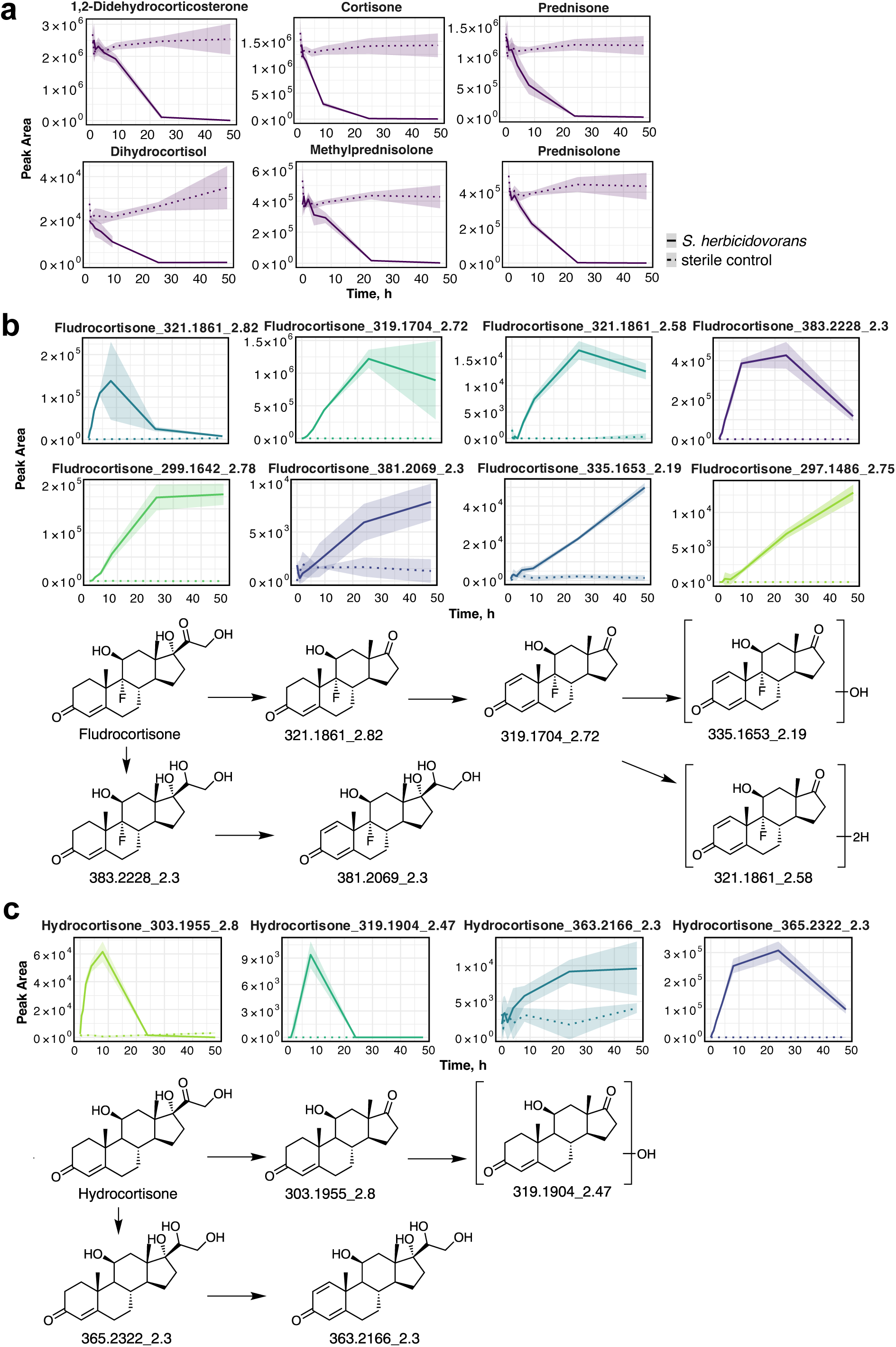
Corticosteroid biotransformation in *S. herbicidovorans*. **a,** Degradation curves of additionally tested corticosteroids upon incubation with *S. herbicidovorans* (n = 4 replicates; shaded area, mean ± SD). **b-c,** *S. herbicidovorans*-produced metabolites of fludrocortisone and hydrocortisone and proposed biotransformation pathways highlighting selected metabolites. Despite similar chemical structure, more metabolites with intact steroid core were detected for fludrocortisone, suggesting that fluoro-group decreases accessibility for biotransformation.

**Supplementary Fig. 5.**
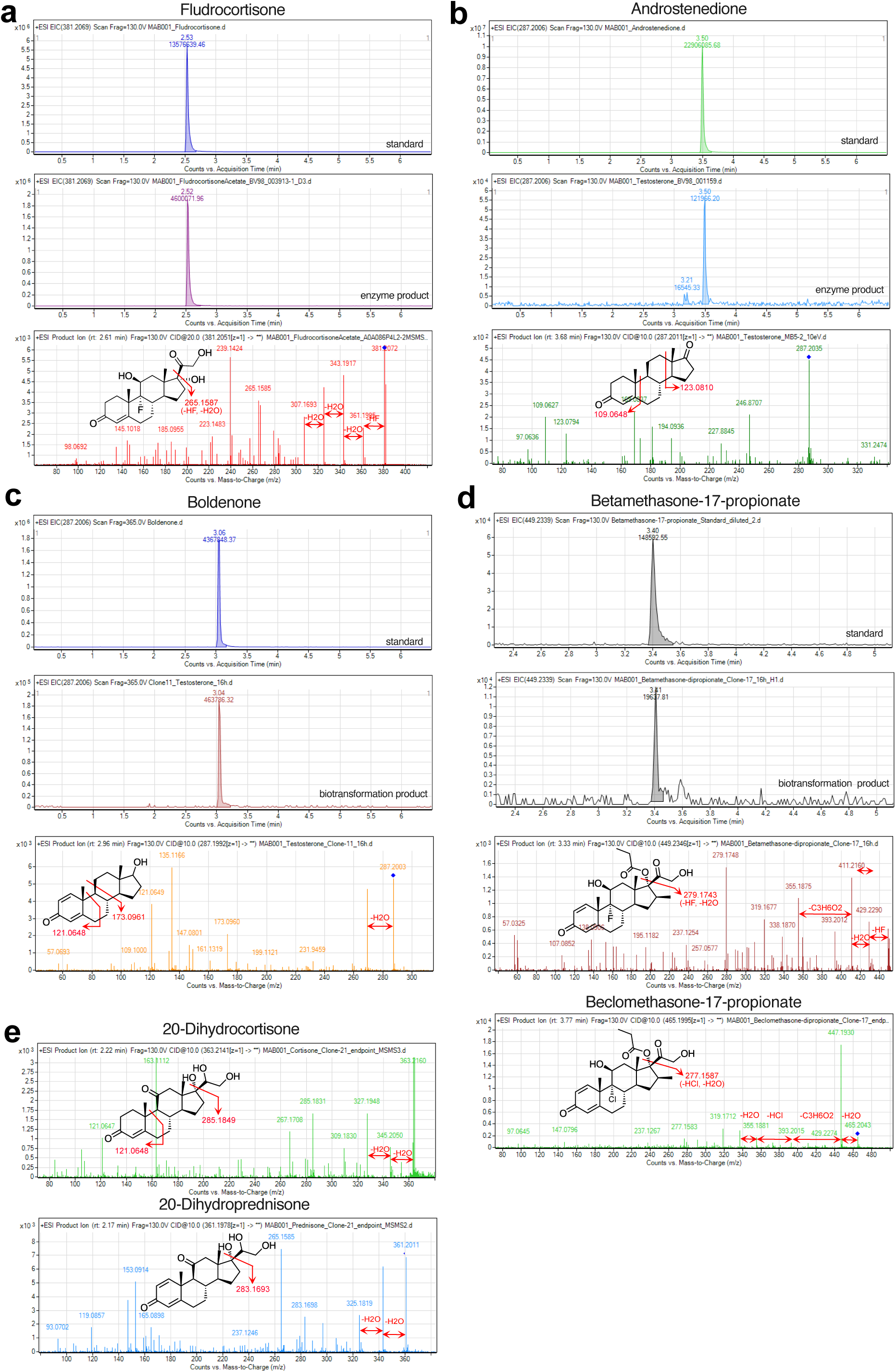
Steroid biotransformation products by *S. herbicidovorans* enzymes. **a,** EIC of fludrocortisone standard, and EIC and MS/MS (20eV) fragmentation of the reaction product of fludrocortisone acetate hydrolase BV98_003913. **b,** EIC of androstenedione standard, and EIC and MS/MS (10eV) fragmentation of the reaction product of testosterone oxidoreductase BV98_001159. **c,** EIC of boldenone standard, and EIC and MS/MS (10eV) fragmentation of the product of testosterone 1,2-oxidation by *E. coli* expressing 1,2-dehydrogenase BV98_003248. **d,** EIC of betamethasone-17-propionate standard, and EIC and MS/MS (10eV) fragmentation of the reaction product of *E. coli* expressing hydrolase BV98_003946, and MS/MS (10eV) fragmentation of the same reaction product with beclomethasone-17-propionate. **e,** MS/MS (10eV) fragmentation of 20-dihydrocortisone and 20-dihydroprednisone produced by *E. coli* expressing 20-ketosteroid hydrogenase BV98_003246.

**Supplementary Fig. 6.**
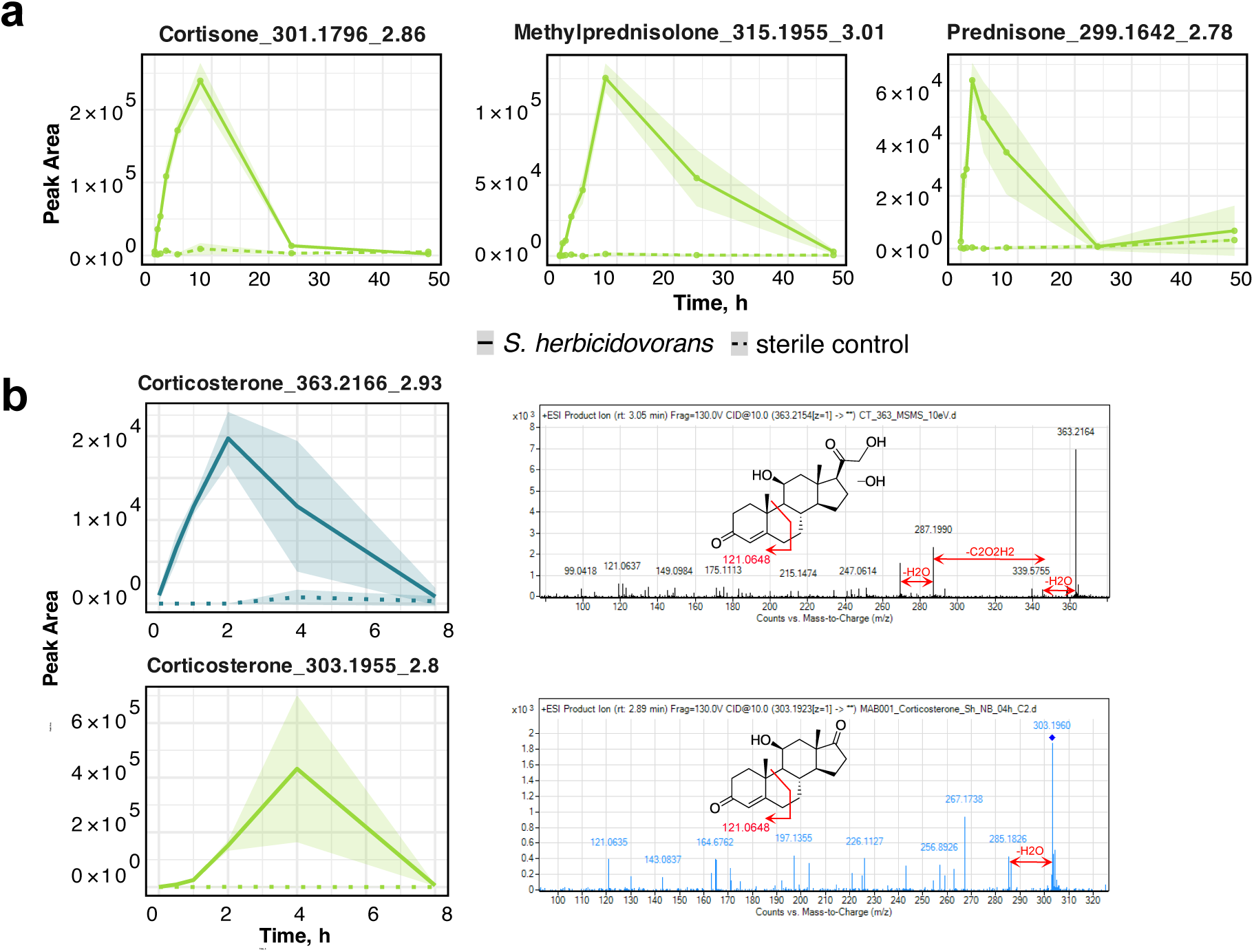
Side-chain cleavage in *S. herbicidovorans*. **a,** Side-chain cleavage of cortisone, methylprednisolone and prednisone in *S. herbicidovorans* leads to the production of 17-ketone metabolites (n = 4 replicates, shaded area, mean ± SD). **b,** Kinetics (n = 4 replicates, shaded area, mean ± SD) and MS/MS (10 eV) for products of corticosterone biotransformation by *S. herbicidovorans* including oxidation/hydroxylation metabolite and side-chain cleavage.

**Supplementary Fig. 7.**
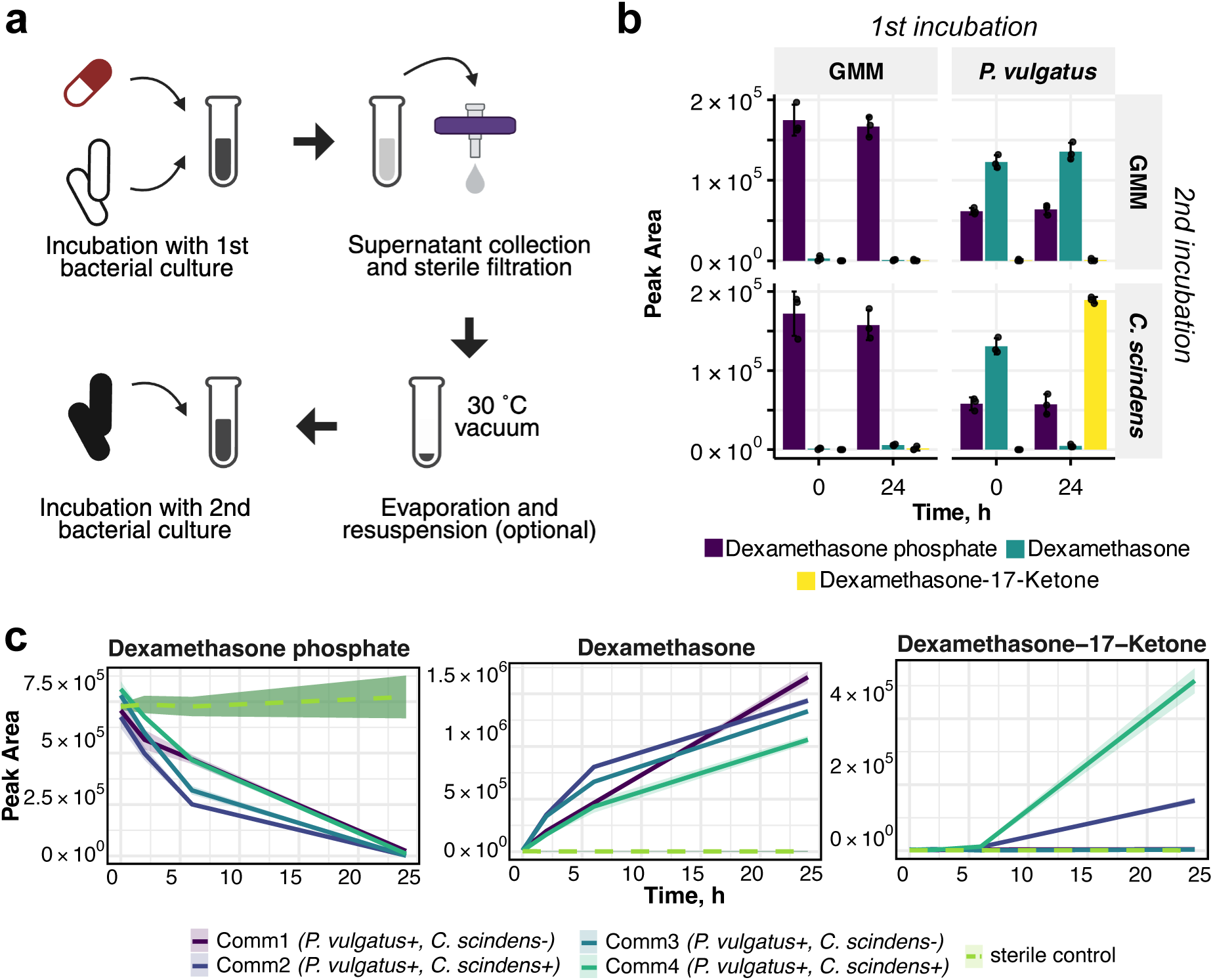
Cross-feeding of dexamethasone phosphate in single species and microbial communities. **a,** Schematic overview of the cross-feeding assay including first incubation, supernatant filtration and evaporation and second incubation. **b,** Cross-feeding of dexamethasone phosphate between gut bacteria *P. vulgatus* and *C. scindens* (n = 3 replicates; first incubation, 48 h; cross-feeding incubation, 24 h; error bars, mean ± SD); GMM is a sterile media control. **c,** Dexamethasone phosphate conversion by four human-stool derived bacterial communities (n = 4 replicates per community; shaded area, mean ± SD).

**Supplementary Fig. 8.**
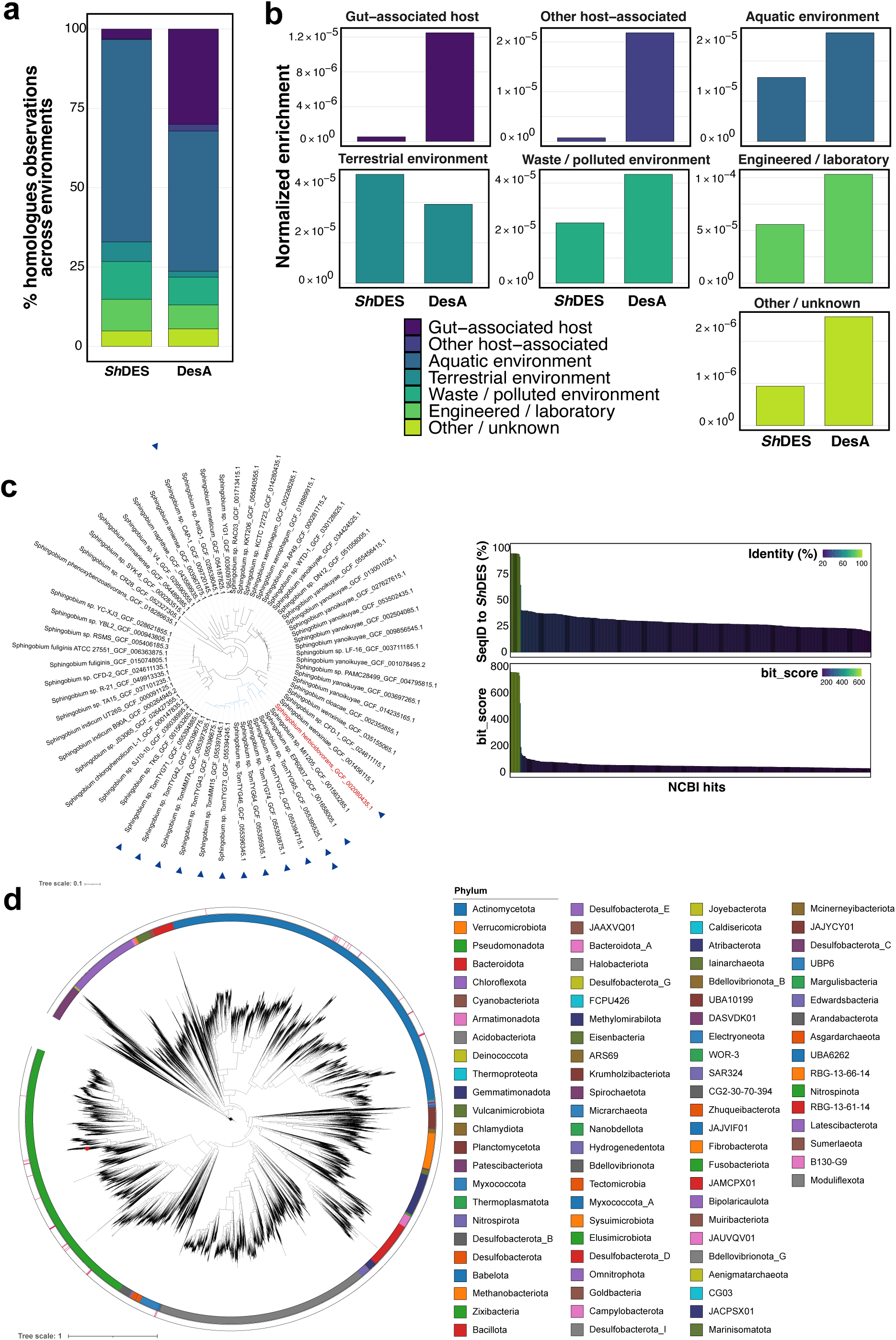
Profiling *Sh*DES desmolase homologues in other species and MGnify metagenomes. **a,** Relative prevalence of detected *Sh*DES and DesA homologues across environments, expressed as the percentage of homologue observations assigned to each biome group. **b,** Enrichment of *Sh*DES and DesA homologues across environments, calculated as the number of homologue observations in a biome group divided by the total number of MGnify observations associated with that biome group. **c,** Phylogenetic tree of *Sphingobium* species with *Sh*DES homologues mapped onto it; tips are labeled by species and RefSeq assembly accession (GCF), and *S. herbicidovorans* is shown in red (*left*), and per-hit sequence identity to *Sh*DES (SeqID, %) and alignment bit-score for NCBI BLAST hits (*right*). Homologues with SeqID > 50% are marked as blue triangles on the Phylogenetic tree. **d,** *Sh*DES homologues mapped onto a soil metagenome-assembled genome (MAG) collection; the outer ring is colored by phylum (GTDB taxonomy), *Sh*DES is highlighted as red dot, and genomes encoding a *Sh*DES homologue are highlighted with red annotation lines.

**Supplementary Fig. 9.**
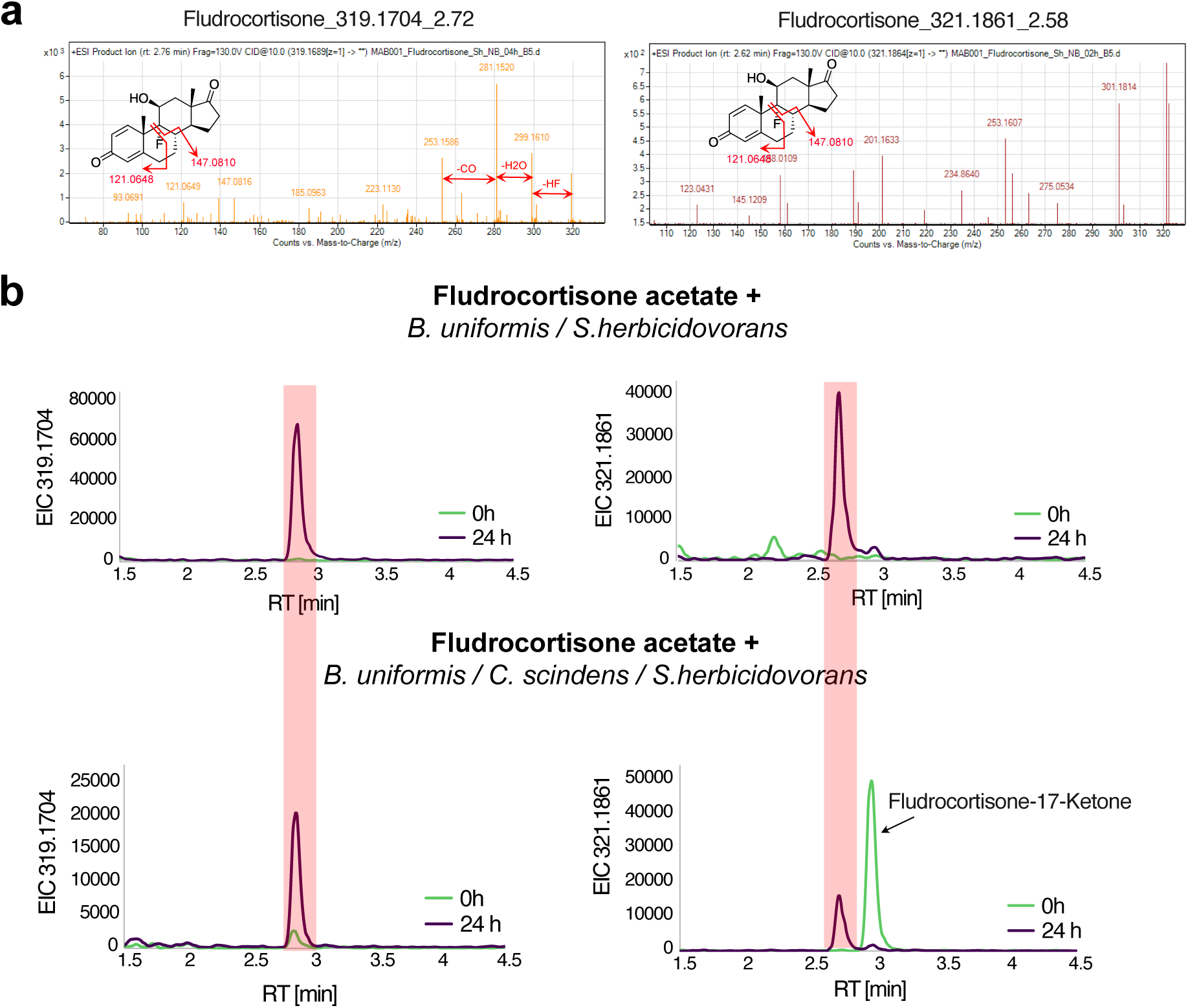
Side-chain cleavage products of fludrocortisone biotransformation. **a,** MS/MS (10eV) fragmentation of the side-chain cleavage products by *S. herbicidovorans*. **b,** EIC of alternative to fludrocortisone-17-ketone side-chain cleavage metabolites produced by *S. herbicidovorans* from fludrocortisone acetate pre-incubated with either *B. uniformis (top)* or *B. uniformis* and *C. scindens,* consecutively *(bottom)*.

